# Uncovering internal states with a robust shared-state multi-neuron GLM-HMM framework

**DOI:** 10.64898/2026.06.27.734988

**Authors:** Aamna Lawrence, Eva Yezerets, Patricia H. Janak, Adam S. Charles

## Abstract

Neural systems exhibit multiple firing states that reflect an organism’s internal state and modulate the relationship between external environmental stimuli and behavior. Several studies have inferred these latent states by supplementing the traditional hidden Markov Model (HMM) with generalized linear models (GLMs) with non-Poisson behavioral observations. However, understanding the relationship between internal brain states and behavior also requires modeling the neural activity. Nonetheless, fitting multi-neuron GLM-HMMs is non-trivial due to high sparsity, collinearity, and low trial counts in neuronal datasets. Therefore, we built a robust multi-neuron GLM-HMM framework that uncovers latent states from population activity while incorporating the influence of time-stamped task variables and spike histories. To obtain reliable model parameters, we employ a modified expectation-maximization procedure. Specifically, we show that incorporating neuron-adaptive penalization in the maximization step overcomes the covariate co-linearity issues typical of time-stamped events and sparse spiking, yielding stable estimates of Poisson GLM coefficients. Furthermore, we incorporate a trust-region algorithm to ensure stable M-step convergence in the presence of ill-conditioned Hessians that can lead to unstable Newton-Raphson updates. We further demonstrate the utility of leave-one-out cross-validation analysis for evaluating model performance on datasets with low trial counts and without breaking their temporal structure. We evaluate our framework on three electrophysiological datasets from primates and rodents as they perform a decision-making task, demonstrate stable model convergence, and discuss the behavioral relevance of the inferred states.

**Author Summary:** Neural systems evolve over time: not only do the individual neurons influence each other across the network, but the network and interconnections themselves change as an animal enters different behaviors (e.g., attentive vs. disengaged) or states (e.g., hungry or tired). Analyzing the neural activity that guides behavior thus must incorporate the time-varying nature of the brain. Recent modeling work has extended the popular Generalized Linear Model, a model that can connect task and behavior to recorded neural action potentials, to incorporate a latent Hidden Markov Model. This extension allows the resulting GLM-HMM to exhibit several different relationships (different GLMs) that are switched between over time to account for the animal’s changing patterns. While GLM-HMMs have been applied extensively on behavioral data (e.g., task choice in a decision making paradigm), neural data is much more difficult due to the smaller sample sizes, sparser activity, and larger parameter space. Our work presents a new fitting approach and best practices to robustly fit GLM-HMMs to neural data. We demonstrate through numerous applications to a variety of neural datasets that by robustly fitting GLM-HMMs to data, we can identify important features of neural activity that let us better understand its relationship to behavior.

## 1 Introduction

Nervous systems are highly dynamic networks with rich underlying processes, often called internal states. These unobserved states modulate how environmental and physiological inputs are integrated in the brain, enabling the organism to generate appropriate behavioral and physiological responses (Karigo and Charles, 2025). With advances in high-throughput behavior-tracking techniques, internal states can be inferred from behavior by clustering similar user-identified animal pose and action features (Mathis et al., 2018; Wu et al., 2020; Pereira et al., 2022). Additionally, progress in data-driven approaches has enabled the use of unsupervised probabilistic methods, such as hidden Markov Models (HMMs), to infer underlying states from behavior. Calhoun et al. (2019) and others (Ashwood et al., 2022; Bolkan et al., 2022; Hulsey et al., 2024; Cuturela et al., 2025; Mohammadi et al., 2025; Li et al., 2023; Chen et al., 2026) have successfully combined HMMs with regression methods, such as generalized linear models (GLMs), to capture varying trial-by-trial behavioral strategies as fluctuating latent states that exhibit distinct relationships with stimuli and other covariates, including internal goals and needs. Such studies have challenged the classical computational models of perceptual decision-making that assume the same trial-based strategy (Green and Swets, 1966; Ratcliff and Rouder, 1998; Ratcliff and McKoon, 2008; Gold and Shadlen, 2007). However, inferred states from behavioral datasets cannot capture the motivational differences between poses or actions that look similar; for instance, mounting behavior could suggest aggression or mating (Karigo et al., 2021). Thus, internal state information from poses and movements must be supplemented with neuronal computations to achieve a comprehensive understanding of complex behaviors.

Prior work suggests that behavioral states are encoded and can thus be predicted from the neuronal population dynamics. For instance, Gründemann et al. (2019) showed that two distinct and antagonistic subpopulations of basal amygdala neurons drive exploratory and defensive behaviors in mice. Similar results have been reported in fish (Marques et al., 2020) and worms (Ji et al., 2021). Furthermore, using artificial stimulation techniques of subpopulations of neurons projecting to multiple brain regions, scientists have successfully induced brain states that influence a range of behavioral features like hunger, thirst, aversion, aggression, courtship, and attention (Aponte et al., 2011; Berrios et al., 2021; Livneh et al., 2017, 2020; Chen et al., 2019; Churgin et al., 2017; Flavell et al., 2013; Ji et al., 2021; Asahina et al., 2014; Anderson, 2016; Clowney et al., 2015; Hindmarsh Sten et al., 2021). Thus, uncovering ensemble activity patterns through unsupervised machine-learning algorithms like GLM-HMMs can help quantify the relationship between underlying neuronal dynamics and behavior.

Escola et al. (2011) first introduced GLM-HMMs for neural activity. However, while behavioral GLM-HMMs have found widespread use in explaining strategy switches across trials, as discussed above, current approaches for leveraging neuronal GLM-HMMs to do the same can be unstable, prohibiting their adoption. Specifically, current fitting methods for GLM-HMMs applied to neuronal data are sensitive to data statistics, parameter configurations, and the amount of data available for model training. These challenges arise as neural data is often degenerate due to sparse or highly correlated activity, has limited trials, and is higher-dimensional.

To remedy the above limitations, we modify the GLM-HMM fitting framework to be more stable under degenerate data and small sample sizes. First, we introduce a model penalization term to handle overfitting and collinear features. Next, we stabilize M-step convergence for ill-conditioned Hessians, and lastly, we assess model fit beyond a conventional train-test split for datasets with low trial counts. We test our model improvements on three electrophysiology datasets in which animals engage in decision-making, demonstrating not only stable fitting but also meaningful latent-state interpretations of behavior.

In this work, we first give an overview of the use of GLMs and HMMs on neuronal datasets in Section 2. Section 3 then provides the mathematical overview of the shared-state population GLM-HMM framework. In Section 4 we present our proposed improvements for a robust GLM-HMM framework. In Section 5, we then investigate the application of our improved shared-state GLM-HMM fitting on three electrophysiological datasets from 1) rodent medial prefrontal cortex (Peyrache et al., 2009, 2018), 2) primate premotor cortex (Lawlor et al., 2018; Perich et al., 2018a), and 3) rodent anterior insula. We assess our model’s fit and compare it with the previous GLM-HMM framework (Escola et al., 2011) across all datasets to highlight the benefits of each design choice. We also discuss meaningful state interpretations in relation to behavior. Lastly, in Section 6, we conclude with a brief summary of the methodological advances introduced by our framework and its applications in neuroscience.

## 2 Background

The GLM-HMM model, as per its name, comprises two main components: a Hidden Markov Model and a Generalized Linear Model. Here we review these components and define notations for the rest of the paper.

### Generalized Linear Models

Generalized linear models allow for linear predictors to be defined under non-Gaussian models. In general, these models define a likelihood distribution of the data (indexed by *t*, which will later correspond to time) ***y***_*t*_ as *p*(***y***_*t*_|***x***_*t*_; *θ*), where ***x***_*t*_ is a vector of regressors and ***θ*** is the set of model parameters, and the expectation of ***y***_*t*_ is a linear function of ***z***_*t*_:

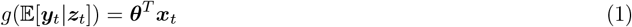

where here *g*(·) is referred to as the link function. Thus the mean of the likelihood is a linear-nonlinear function of the regressors ***x***_*t*_.

Here we are specifically interested in fitting models to spike data, describing spike counts as a function of task variables and spike history. In this case, the likelihood function over the emission matrices follows a Poisson distribution, such that the probability of observing *y*_*t*_ spikes from a given neuron at time *t* is given by

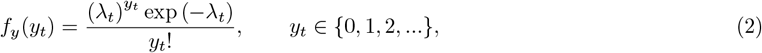

where *λ*_*t*_ is the firing rate of the neuron in time-bin *t*. By the GLM assumption we have, taking the typical log(·) link function

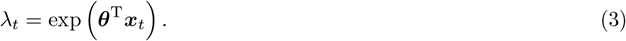

Here, ***θ*** ∈ ℝ^*D*^ represents the weight vector and ***x***_*t*_ ∈ ℝ^*D*^ is the covariate vector at time *t*. ***x***_*t*_ includes variables such as the task variables and spike history and potentially spiking activity of other neurons (Paninski, 2004). The spike history component consists of spike counts *τ* steps prior to *t*: **y**_*t*−*τ*:*t*−1_ = (*y*_*t*−1_,…, *y*_*t*−*τ*_).

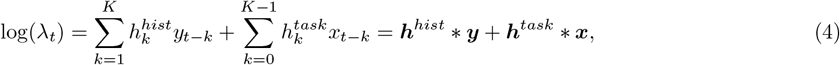

where ***h***^*hist*^ and ***h***^*task*^ are length-*K* filters that dictate how the past history and task variables, respectively, influence the current spike rate. Fitting the filters as a function of time can be sensitive to noise and available data, so often a prior is used wherein the filters are expanded into a set of *L* basis elements **Φ** = [***ϕ***_1_, ***ϕ***_2_, …, ***ϕ***_*L*_] ∈ ℝ^*K×L*^ as

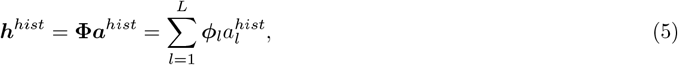

for ***h***^*hist*^ and similarly

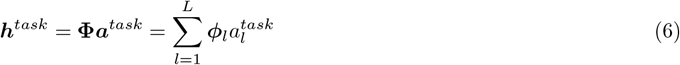

for ***h***^*task*^. Here the coefficients ***a***^*hist*^ and ***a***^*task*^ represent how much of each basis element is used to construct each filter, and are the free parameters in the GLM that are to be fit. With estimates of ***a***^*hist*^ and ***a***^*task*^, we can reconstruct the actual filters ***h***^*hist*^ and ***h***^*task*^, respectively as ***h***^*hist*^ = **Φ*a***^*hist*^ and ***h***^*task*^ = **Φ*a***^*task*^. With these definitions we can rewrite the GLM expression for the log-expectation in terms of ***a***^*hist*^ and ***a***^*task*^ as

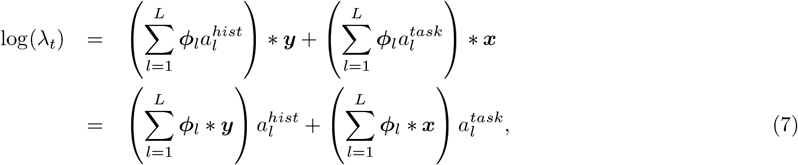

where we can reorganize the terms due to the linearity of convolution. This expression means that the regressor for each basis of the basis expansion of the kernels ***h*** are the convolution of the corresponding time-series (i.e., the spike history or the task inputs/outputs) with the corresponding basis elements. Following earlier (Park et al., 2014) in GLMs for neural activity, we use as the basis vectors ***ϕ***_*l*_ for *l* = 1, 2, …, *L*, a set of raised cosine basis functions given by

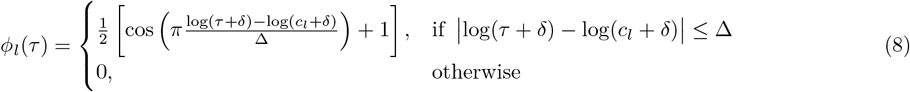

where *δ* is the small positive offset to prevent divergence near zero, *c*_*l*_ is the center of the *l*^*th*^ basis function and Δ is the non-linear spacing between the bases. Optimized versions of Poisson GLMs have also been proposed (Medvedeva et al., 2025) that overcome binning of spike data to construct efficient design matrices and enable precise functional connectivity inference using Laguerre polynomials as basis functions.

While the GLM described above is a good *local* model for any particular time-point, we note that the specific interactions learned in ***θ*** might change over time. These changes can be a consequence of learning, adaptation, neuromodulation, or context changes in the brain (among other possibilities). Thus neural time-series might be better represented by a set of *S* GLMs where each *y*_*t*_ is well represented by one of the *S* GLMs for each *t*. We can denote these different GLMs by indexing the model parameters, i.e., ***θ***_*s*_ for *s* = 1, 2, …, *S*. However, we do not know *a priori* which GLM best describes each data point (i.e., which state *s* is active at a given time *t*). Thus, the GLM-HMM uses a hierarchical model to overlay a Hidden Markov Model over the state *z*_*t*_ at each time point *t*, where *z*_*t*_ can take one of the *S* discrete states {1, 2,… *S*}.

### Hidden Markov Models

HMMs (Rabiner, 1989) build on the assumption that the state at any time point *t*, denoted *z*_*t*_, is drawn from {1, 2, ….*S*}, and extends a *state transition* assumption that connects the state at any time point to the preceding state. This forms a consistency of states: a state is more consistent if the probability that *z*_*t*_ = *z*_*t*−1_ is high. First, we note that the Markovian assumption of the HMM means that we can write the conditional probability of the state at time *t* as only a function of the previous state; i.e.,

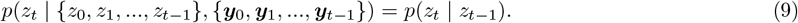

Thus the current state only depends on the state at time *t* − 1 without any impact of the emissions. Next, we define the transition probabilities via the transition matrix ***A*** ∈ ℝ^*S×S*^, where

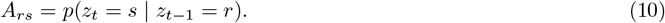

Each entry 0 ≤ *A*_*rs*_ ≤ 1 captures the probability of transitioning from state *s* (the column) to state *r* (the row), and therefore the diagonal *A*_*rr*_ entries are the “stickiness” of the model, or the probability of staying within a given state *r*. ***A*** is homogeneous in that it is assumed constant for all time points. Traditionally, the probability of observing *y*_*t*_ only depends on the current state *z*_*t*_ and not on past states or emission sequences:

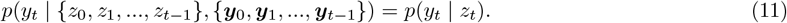

Thus, the full log-likelihood of the model over all the sequence of states is given by

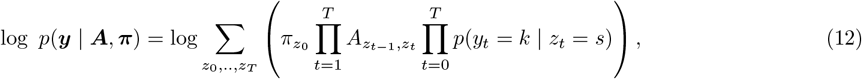

where the system has *K* discrete emission classes {0, 1, 2, ….*K*}, and ***π*** is the initial state probability distribution vector. These model parameters are learned from the data by maximizing Equation (12).

### GLM-HMMs

In combining the GLM and HMM models (introduced first by Bengio and Frasconi (1995)), the main modeling specification is that each distribution *p*(*y*_*t*_ = *k* | *z*_*t*_ = *s*) is defined as a GLM with parameters indexed by the state: ***θ*** → ***θ***_*s*_. In the case of behavioral models, this is, e.g., the logistic distribution (e.g., Calhoun et al. (2019)). Here we follow the above Poisson GLM with the log link function (Simoncelli et al., 2004; Paninski, 2004; Truccolo et al., 2005). This means that each state is characterized by its own set of filters: a per-state spike-history filter, a task-variable filter, etc. as described above. Thus the model can capture different modes of the network, such as if a given task variable sometimes excites, and sometimes inhibits, the same neuron. Such a multi-model response (potentially dependent on contextual or neural cues not observed in the experiment) cannot be captured with a single GLM.

## 3 Shared-state multi-neuron GLM-HMM model

The HMM allows us to find hidden patterns in simultaneously recorded neuronal data, while the GLM provides a way of integrating the task variables and spike history information to understand how the states influence their processing. For ease of notation in the subsequent exposition we define: 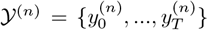 as the full set of activity for neuron *n*, 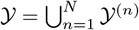 as the full set of neural activity across all neurons, *X* = {***x***_0_, …, ***x***_*T*_} as the full set of inputs across all time, *Ƶ* ={*z*_0_, …, *z*_*T*_} as the set of all latent states across time, and **Θ** = ***θ***_1_, …, ***θ***_*S*_ be the full set of GLM parameter sets across all states.

One additional assumption that can be made is that all simultaneously recorded neurons share a state; i.e., if we assume *N* simultaneously recorded neurons whose spikes are recorded at *T* time points, the probability of observing the ensemble at a given state *z*_*t*_ and the predictors *X* across all time and neurons is,

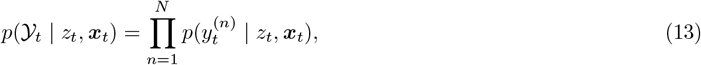

such that each cell’s emission follows a Poisson Distribution given by Equation (2). Thus, the total log-likelihood of the model across all neurons, states and time bins becomes,

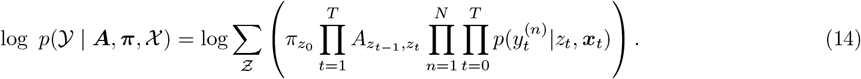

### Baum-Welch Expectation-Maximization algorithm

To fit the model parameters, we would like to maximize the marginal likelihood function in Equation (14). However, this marginal likelihood is intractable to optimize, preventing direct inference. Instead, as with prior work, we turn to the standard Expectation-Maximization (EM) algorithm. Specifically, we use the iterative Baum-Welch EM algorithm (Dempster et al., 1977) to fit the model parameters and optimize the data likelihood. EM proceeds in two steps: During the E-step (Expectation), the posterior probabilities of the latent states are computed given the data and parameters, while in the M-step (Maximization), the next set of parameters is computed by minimizing the negative log-likelihood with respect to the current parameters, using the posterior probability distributions from the E step. These steps are iterated over until a convergence criterion is met.

For ease of notation in the subsequent exposition, for each *X, Ƶ, Y*, and ^(*n*)^ we use the subscripts *< t*, ≤ *t, > t*, ≥ *t*, and /= *t* to indicate the subset of that includes members of the set with index satisfying the subscript’s inequality. Mathematically we can write

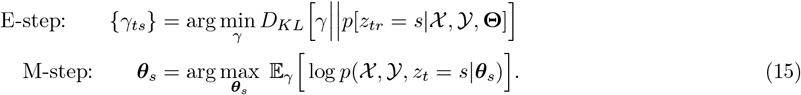

In these equations *γ*_*ts*_ represents the posterior of the active state at time *t* being *s* (i.e., conditioned on the data including the task time series ***x***_*t*_ and neural activity *y*_*t*_, and *D*_*KL*_(*p*||*q*) is the KL-divergence between two distributions.

#### E-step

The E-step in the EM algorithm infers the probability of state membership at each time-point with old model parameters using the forward-backward algorithm (Baum et al., 1970). Effectively the approach approximates the posterior *γ*_*tr*_ = *p*(*z*_*t*_ = *r*|*Y, X*, **Θ**).

To compute *γ*_*tr*_, we note that the marginal posterior probability of being in state *r* at time *t* can be expressed in a form that separates the conditional information from the past and future time-stems. Specifically,

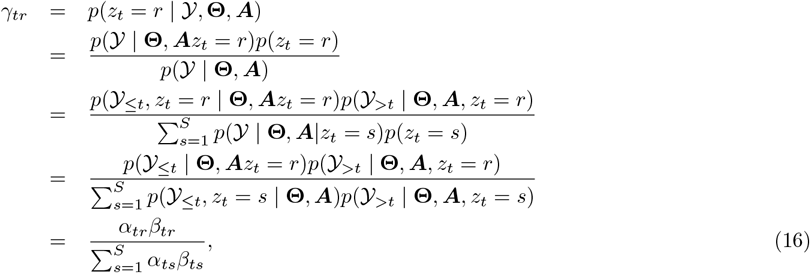

where *α*_*tr*_ and *β*_*tr*_ are the forward and backwards probabilities, respectively, defined as

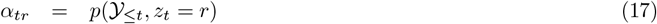

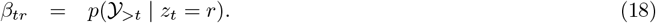

These probabilities are convenient as they can be recursively computed in an efficient manner. For the forward probability, *α*_*t*_(*s*), we can write the distribution as a marginal over all past states,

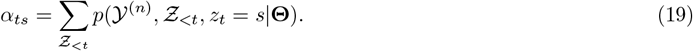

At *t* = 0, this probability is simply

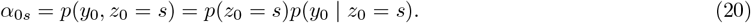

For *t >* 0, we can use the Markov property of the state transition in *z*_*t*_ to write

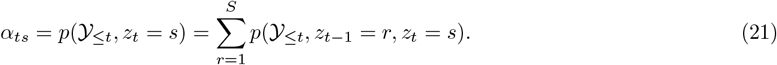

Using the chain rule and the Markovian property from Equations. (9) and (11) in Equation. (21) yields

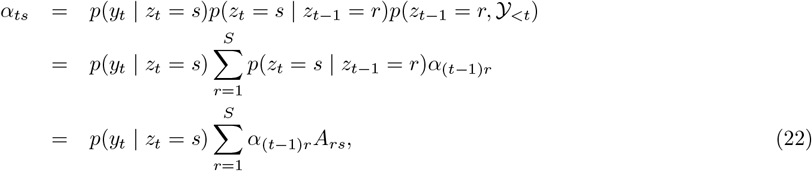

meaning that the current probability of the emission and state depends only on the past distribution of the emissions passed through the HMM state transition matrix.

For the backward probabilities we can similarly express the distribution as the marginal over the next state

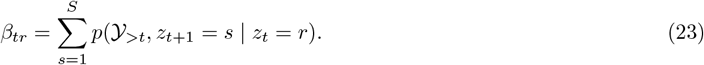

After applying chain rule and using the Markovian property from Equation (11),

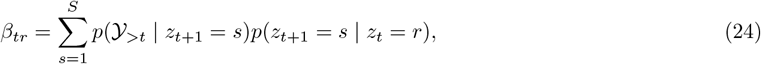

such that,

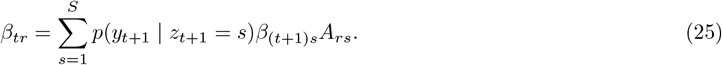

Together, *α*_*tr*_ and *β*_*tr*_ can then be used to compute the esitmates *γ*_*tr*_ as in Equation (16).

#### M-step

In the M-step we use the estimated state probabilities from the E-step in order to update the model parameters: the transition matrix ***A*** and the GLM weights 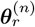 for each state *r* = 1, …, *S* and each neuron *n* = 1, …, *N*. The pairwise posterior probability for all neurons at time *t* is given as,

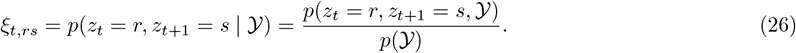

By expanding the numerator and applying the chain rule,

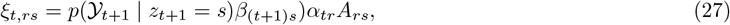

or,

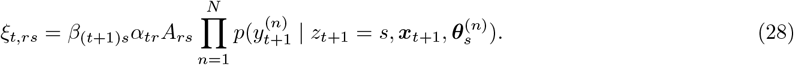

The new transition matrix for the next EM iteration is

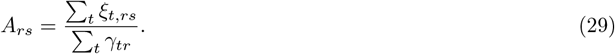

For each state *r* and neuron *n*, the GLM weights 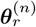 are updated by maximizing the weighted Poisson log-likelihood:

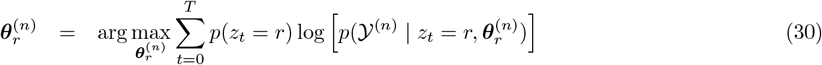

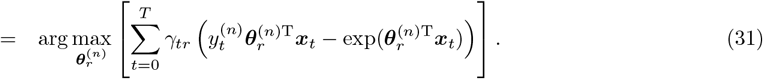

To prevent underflow due to the multiplication of probabilities, Equations (22), (25), (16), and (28) are transformed to the log scale and rescaled for numerical stability (see Supplement).

## 4 Robust GLM-HMM framework

Our robust GMM-HMM fitting framework aims to 1) improve robustness to variability in neuronal activity levels, 2) decrease the sensitivity to the expected ill-conditioned data Hessians, and 3) enable proper model fitting for data-limited experiments.

### Model initialization with neuron-adaptive regularization

Most electrophysiology studies investigate neuronal responses aligned to discrete time-stamped events. To fit a GLM to neuron datasets, the event predictors are binned at millisecond resolution and, to generate the per-basis coefficient regressors as per Equation (7), often convolved with a set of basis functions. The resulting covariate matrices, unfortunately, are typically highly correlated and sparse. Additionally, sparsely firing neurons may be active around certain isolated task events, which can lead to overfitting and the assignment of biologically implausible stimulus coefficients. In contrast, behavioral GLMs often have more well-behaved, continuously varying covariates, which generally produce better-conditioned design matrices and are less susceptible to coefficient inflation.

In the GLM-HMM framework, the issue of poor covariate conditioning is particularly relevant in the M-step, where each neuron has its own state-specific GLM. As a non-convex optimization, parameter initialization can have a strong impact on the estimation stability, especially under the neuron-by-neuron firing rate variability. We thus sought to improve estimation stability while accounting for neuron-specific firing statistics by initializing the GLM weights via a per-neuron, single-state initial parameter fitting. This procedure follows prior initialization approaches in the GLM-HMM literature (Ashwood et al., 2022) with the exception that we introduce a ridge-regularizion term via a Gaussian prior over the weights to induce stable initialization. Mathematically, our modification changes Equation (31) to minimize the log-posterior (including the log-prior as the regularization term) probability as

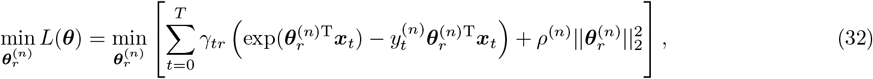

where *ρ*^(*n*)^ is the regularization strength for the *n*^*th*^ neuron, which was solved using the trust-region method (see below). Ashwood et al. (2022) noted that the parameter fits across states can be highly similar, which we also found when including the regularization. We thus also used (as in Ashwood et al. (2022)) an independent Gaussian perturbation scaled to 25% of the norm of the baseline solution to break symmetry between states.

During the initialization we also set the regularization parameter for each neuron *ρ*^(*n*)^ by fitting single-state, ridge-regularized GLM with five-fold cross-validation. The neuron-specific ridge penalty selected during initialization is retained and used in M-step optimization, which improves numerical stability and keeps the coefficients well-constrained.

### Robust M-step convergence using the trust-region method

The weights for each neuron *n* in a particular state are updated for each EM iteration by the above optimizing Equation (32) that minimizes the negative log-posterior *L*(***θ***). Optimizing this code function depends on computing the gradient ∇_*θ*_*L*(***θ***) and Hessian 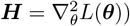, which are expressed as:

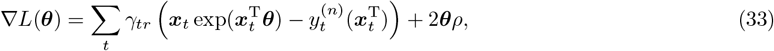

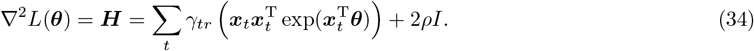

The Newton-Raphson method approximates the optimization function *L*(***θ***) locally about the current estimate of ***θ*** with a quadratic surrogate function. This surrogate allows for an efficient update that accounts for both the gradient and curvature via a Newton iteration as:

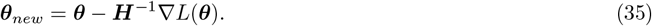

However, if the local curvature is anisotropic, ***H*** can become ill-conditioned, resulting in a large Newton step and unstable weight estimates.

To prevent such large weight updates during optimization, we instead use a trust-region method. At each iteration, a second-order Taylor approximation of the objective function *L*(***θ***) is computed, given as

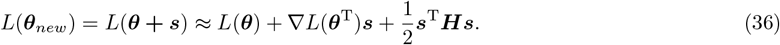

By ensuring that the step, ∥***s***∥_2_ *<*= Δ, where Δ is the radius within which the step is taken, the step is computed as

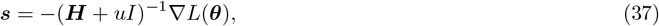

for some *u* ≥ 0 such that || ***s*** || ≤ Δ. If *L*(***θ* + *s***) *> L*(***θ***), the region of trust is shrunk and vice-versa. The trust-region optimization was implemented using MATLAB’s fminunc solver with the trust-region algorithm in the M-step.

### Model fit cross-validation with minimal data

Conventional k-fold cross-validation may not be optimal for trial-based electrophysiology datasets due to two reasons: 1) randomly partitioning temporally-structured observations can disrupt within-trial dependencies that are necessary for GLM-HMM inference, and 2) low-trial-count datasets may not provide enough data for appropriately training the model.

We therefore use a leave-one-trial-out cross-validation procedure for assessing our model. Specifically, if a dataset has *N* trials, we train our model on *N* − 1 trials, test it on the *N* ^*th*^ trial, and repeat until all trials are tested. For computing the log-likelihood of the held-out test trial, we run only the forward pass and use the parameters estimated after training the model on *N* − 1 trials. For each held-out test trial, predictive performance is quantified in bits per second, described by Calhoun et al. (2019) as:

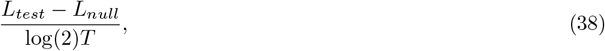

where

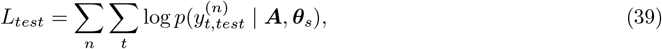

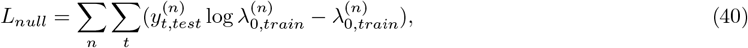

where 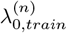 is the mean firing rate of the *n*^*th*^ neuron in the training trials, and *T* is the total trial duration of the held-out test trial. To assess if the predictive performance is statistically significant, Equation (38) is recomputed on the held-out test trials for the model trained on shuffled neuronal spikes per trial.

## 5 Fitting to electrophysiology data

In this section, we apply our framework to three electrophysiology datasets, including neurons with a mean firing rate of ≥ 0.2 Hz across the session, to assess its ability to learn the model parameters appropriately and converge stably.

### Medial prefrontal cortex during a contingency task

We first applied our methods to a medial prefrontal cortex (mPFC) dataset in which freely moving rats performed an attentional shift task (Peyrache et al., 2009, 2018). Each trial started with the rat in the root of a Y-maze. According to the spatial contingency rule, one of the choice arms (the right arm in the dataset) was rewarded. We trained our model to a session where the animal performed 22 trials (13 correct and 9 incorrect) and had 31 neurons recorded simultaneously. To prepare the predictor matrix, trials were concatenated across the session and binned into 50-millisecond time bins. Timestamps for trial start and end for choice and outcome encoding, respectively, along with the spike history were used predictors and convolved with 5 raised-cosine basis functions with a time support of 2.5 seconds. No position information was given to the model. Since the true number of hidden states was unknown, we considered *S* ∈ {3, 4, 5}. While the negative log-likelihood of our model decreased as *S* increased, the state identities became less clearly defined (Supp. Fig 1). Therefore, we decided to select the 4-state model.

Our model revealed that the recruited states sequences differed between the correct and incorrect trial types (Fig 1). Importantly, at trial start and end, when the animal attended to the contingency, went to the correct arm, and received the reward, state 3 was on, whereas when the animal went to the incorrect arm, state 2 was on. To gain further insight into population dynamics, we investigated the population-averaged kernels at trial start for states 2 and 3. The kernel modulation was higher and positive in state 3 than in state 2 (see the insets in Fig 1), highlighting active engagement of the mPFC at trial start. These results corroborate prior studies implicating the mPFC in the representation of reward-associated memories during task performance (Birrell and Brown, 2000; Floresco et al., 1999; Jones and Wilson, 2005).

**Figure 1.**
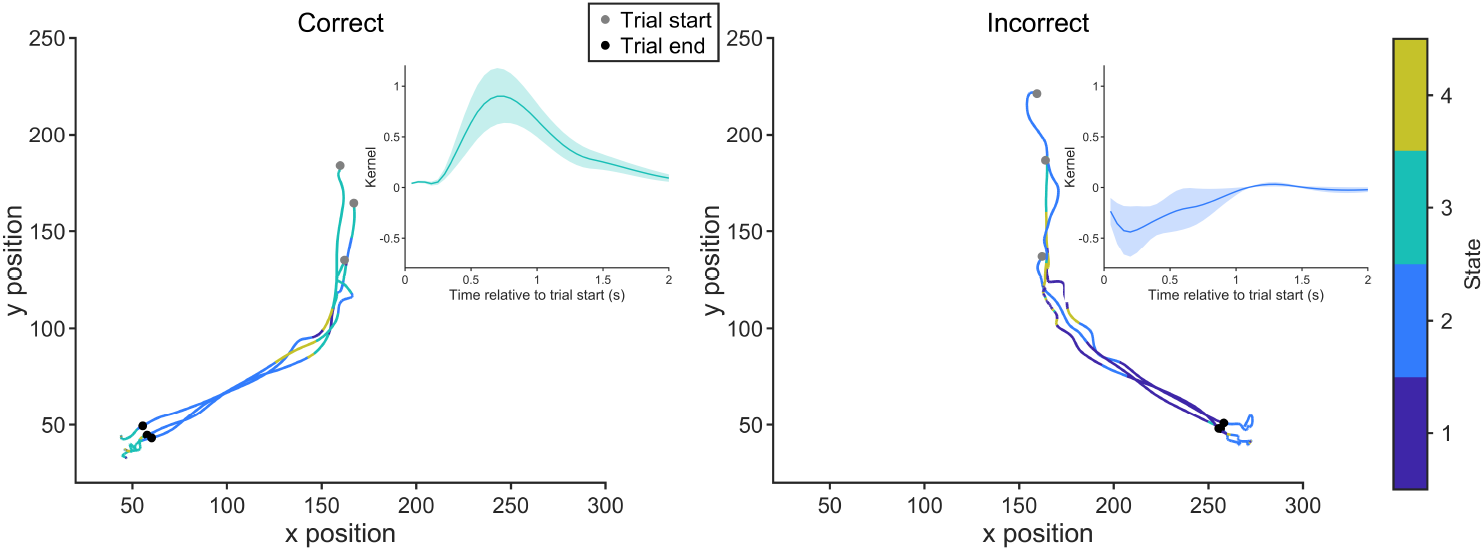
Attention-contingent state recruitment in the mPFC. Three example correct (turn right) and incorrect (turn left) animal position trajectories color-coded by the highest probability state in an attentional shift task. Insets show population-averaged kernel deviations for states 3 and 2, associated with correct and incorrect trial starts, respectively (Mean±SEM).

Additionally, our model showed an appropriate decrease in the negative log-likelihood and squared error in the model parameters across the EM iterations, highlighting stable convergence (Fig 2A,B). Furthermore, the average per-neuron correlation between the GLM weights obtained from the M-step were minimally correlated (Fig 2C), proving that the states were non-redundant. To establish that our model could extract meaningful information when trained on such low trial counts, we performed a held-out trial analysis where we trained the model on 21 trials and used the model parameters to estimate the log-likelihood on the 22^nd^ held-out trial (Fig 2D). Our model showed a significantly higher held-out likelihood on the test trial than when it was trained on shuffled spike data (Wilcoxon signed-rank test).

**Figure 2.**
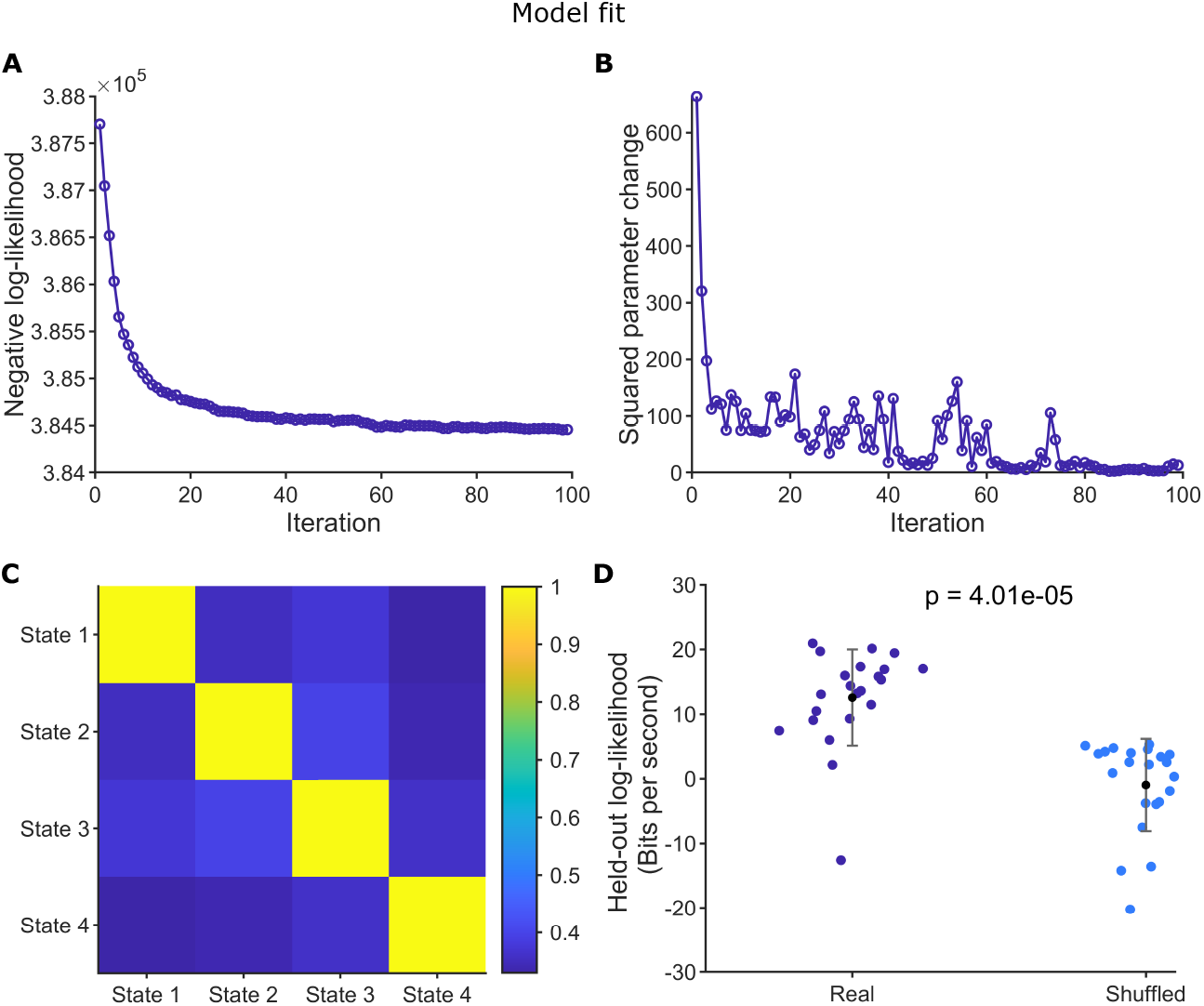
Shared 4-state GLM-HMM validation and convergence evaluation for rodent mPFC activity. (A) Negative log-likelihood across EM iterations. (B) Change in model parameters (***A*, Θ**) across EM iterations. (C) Mean GLM weight correlation between states per neuron. (D) Held-out log-likelihood for model trained real and shuffled spikes from 21 trials and tested on 22^nd^ trial (Wilcoxon signed-rank test). Each dot represents the held out trial.

The initial model weights for each neuron were calculated by fitting a Poisson GLM with a ridge penalty with five-fold cross-validation. Subsequently, this neuron-adaptive regularization was used in the M-step to ensure that the weights remained within biological limits. We compared our model initialization with two alternatives: (1) initialization using an unregularized GLM, and (2) random initialization without GLM-based fitting. Fig 3A shows the distribution of GLM weights for each method for the final EM iteration. The unregularized GLM initialization produced heavy-tailed weight distributions, indicating that a subset of coefficients reached extreme magnitudes due to overfitting. The random initialization coefficients, while not as extreme as those of the unregularized GLM, still contained a subset of high, biologically implausible coefficients. In contrast, our ridge-regularized initialization resulted in well-bound weight distributions.

**Figure 3.**
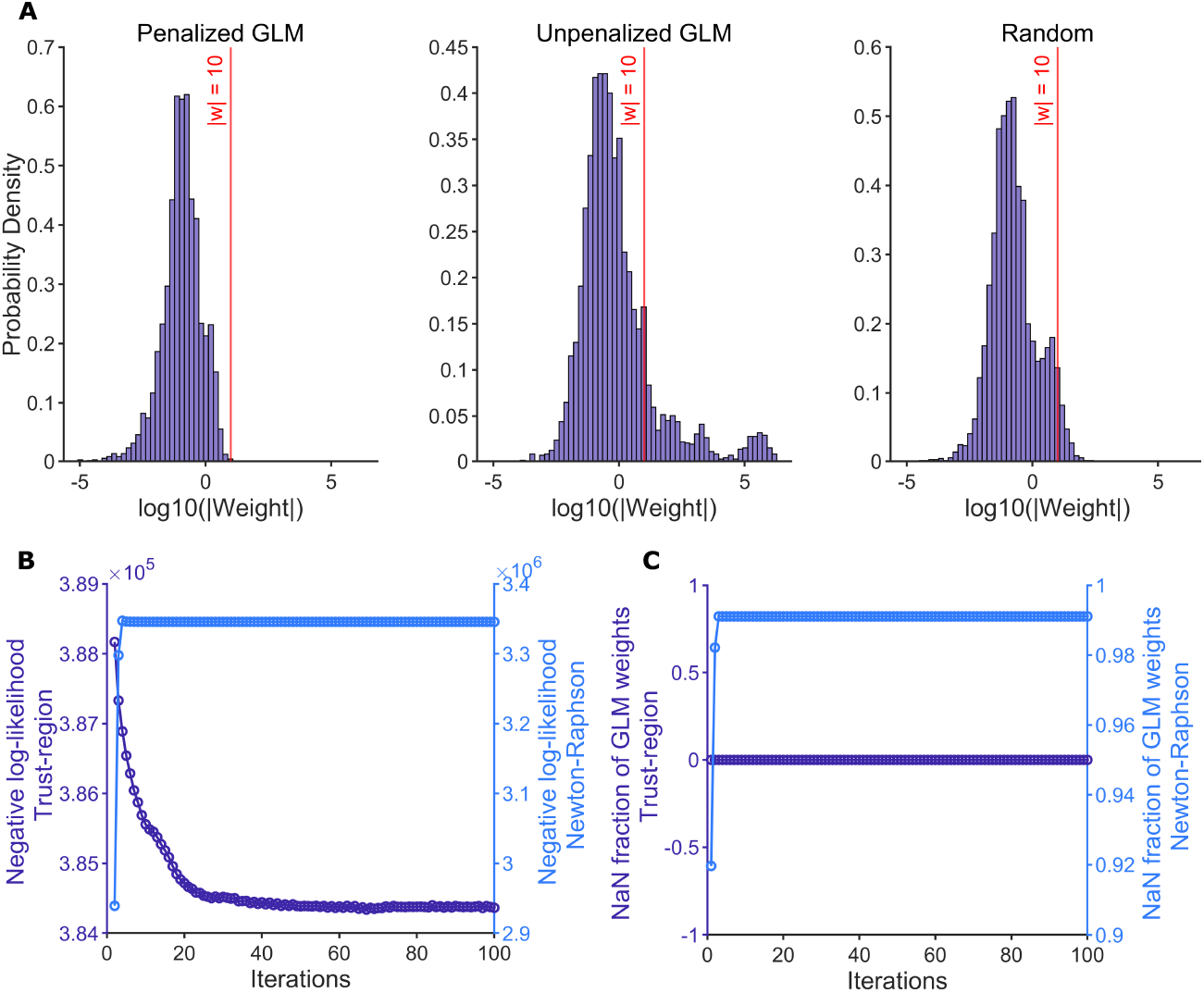
Comparison of initialization and optimization strategies for GLM-HMM fitted on mPFC activity. (A) Probability density distribution of final absolute GLM weights in log10 scale across all neurons using penalized GLM, unpenalized GLM, and random initializations. (B) Negative log-likelihood comparison across iterations when using trust-region and Newton-Raphson methods for M-step convergence. (C) Fraction of non-finite GLM weights across EM iterations when using trust-region and Newton-Raphson methods in the M-step.

To evaluate the convergence of our shared 4-state model, we compared optimization using the trust-region method in the M-step to a pure Newton-Raphson approach. We tracked the negative log-likelihoods and GLM weights for every EM iteration. The negative log-likelihood exponentially decreased when the model used trust-region method unlike the Newton-Rapson method, showing that Newton-Raphson is not the best way to optimize the GLM weights in the M-step for sparse electrophysiology datasets (Fig 3B). Additionally, the Newton-Raphson updates led to numerical instability: 99% of the weights become non-finite after the second EM iteration. The trust-region method, by comparison, maintained fully finite parameter estimates for all iterations (Fig 3C).

### Premotor cortex during reach task

We next applied our methods to a premotor cortex dataset in which monkeys were trained to perform a sequential reach task to move the computer cursor to a visual cue specifying the target location. Four sequential correct reaches resulted in liquid reward delivery (Lawlor et al., 2018; Perich et al., 2018a). The analyzed dataset had 94 simultaneously recorded neurons and a total of 496 reaches. To prepare the predictor matrix, target reach epochs were concatenated across the session and binned into 10-millisecond time bins. Timestamps for each target onset event (approximated) and spike history were used as predictors. These events were convolved with 5 raised cosine basis functions with a time support of 0.4 seconds. We chose a 2-state model since a 3-state model exhibited strong correlations among GLM weights per neuron (Supp. Fig 2).

Our two-state model revealed that for the boundary targets (targets 1 and 4, corresponding to the movement sequence initialization and termination), state 1 was consistently recruited, whereas for intermediate targets (targets 2 and 3, corresponding to ongoing sequence execution), state 2 was recruited (Fig 4A,B). Furthermore, the population-averaged spike history kernel was significantly different for the two states (Fig 4C), which suggests that the premotor cortex dynamics differ within a trial. The consistent state recruitment across rewarded trials also shows that our model captured trial-by-trial variability in population neuronal activity without explicitly implementing dynamic time warping as previously done by Lawlor et al. (2018).

**Figure 4.**
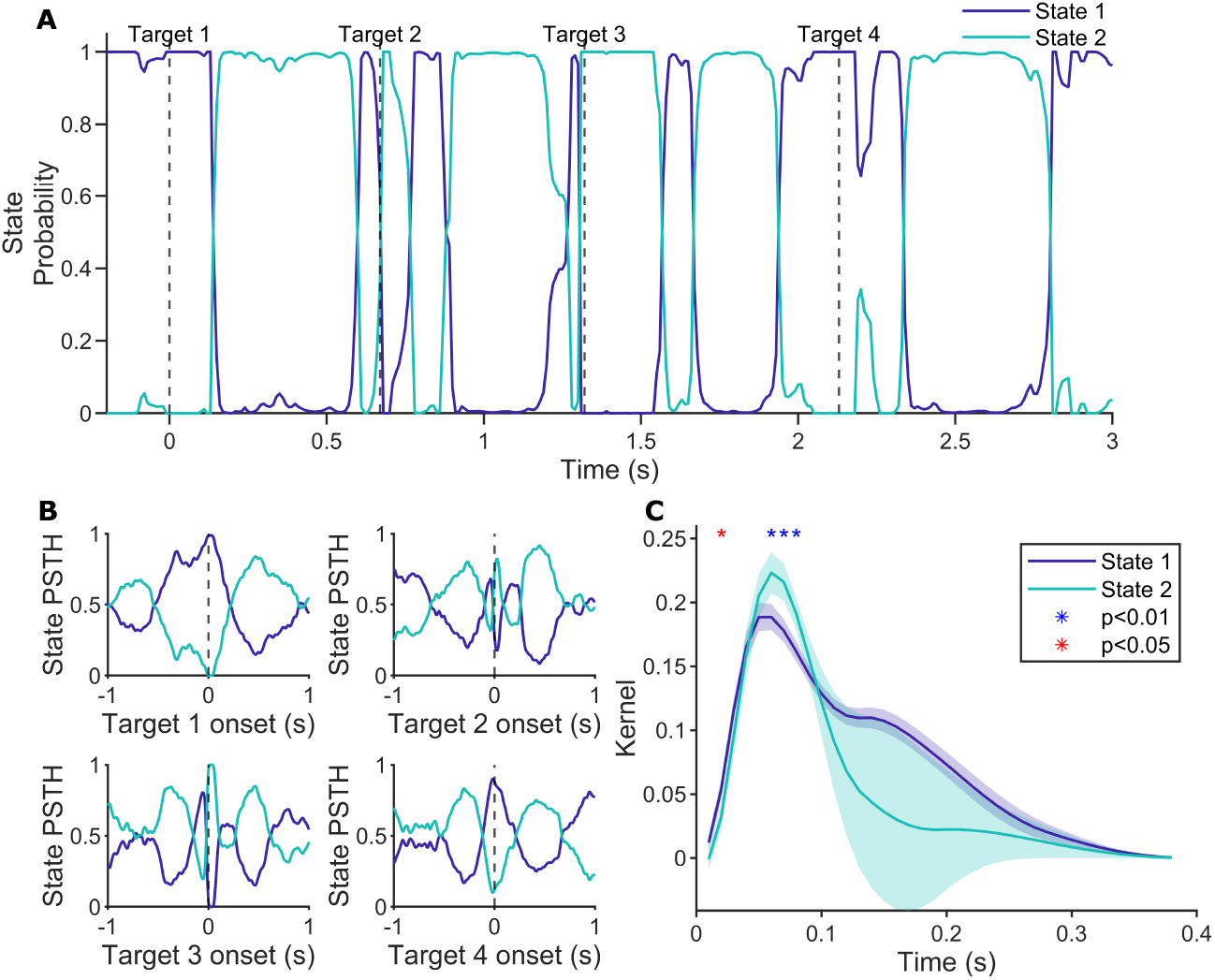
Consistent recruitment of distinct states for intermediate and peripheral targets across trials in a sequential target reach task. (A) Posterior probabilities for an example trial. (B) Fraction of trials assigned to each state ± 1s relative target onset (Gaussian-smoothened over 5 time bins).(C) Population-averaged spike history kernel deviations for states 1 and 2 (Mean±SEM; Wilcoxon signed-rank test).

Similar to the 4-state model trained on the mPFC dataset, our 2-state model showed an appropriate decrease in the negative log-likelihood and squared error in the model parameters across the EM iterations, highlighting stable convergence (Fig 5A, B). Furthermore, the average per-neuron correlation between the GLM weights obtained from the M-step were minimally correlated, showing state non-redundancy (Fig 5C). We also did a held-out trial analysis by training our model on a subset of reaches (133 reaches to target 2). Even with a quarter of the total reaches in the entire dataset, the model trained on real data outperformed the model trained on the shuffled dataset, suggesting that our model could capture the population activity patterns in the premotor cortex with minimal data (Fig 5D; Wilcoxon signed-rank test).

**Figure 5.**
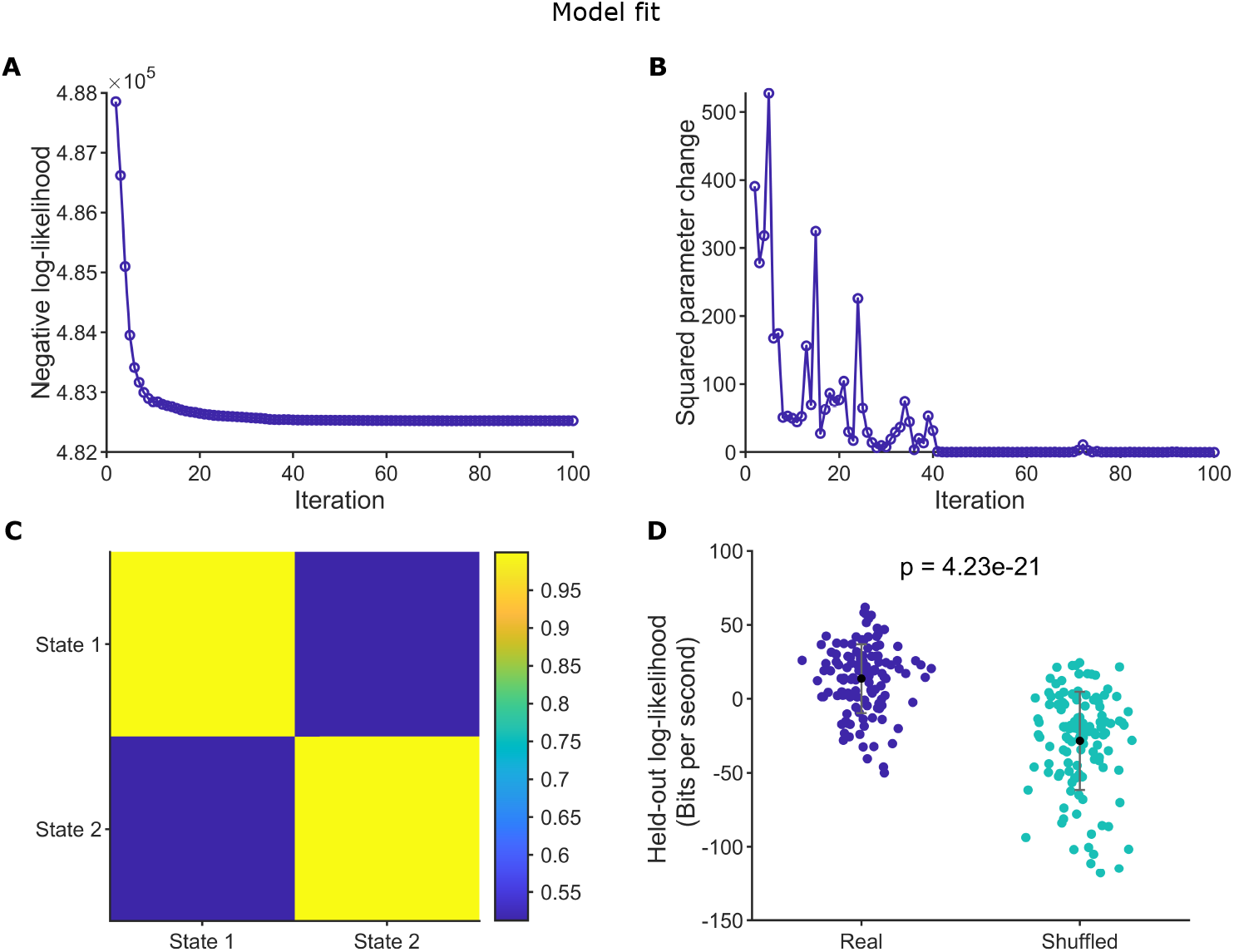
Convergence and validation of the 2-state GLM-HMM trained on premotor cortex data. (A) Negative log-likelihood across EM iterations. (B) Change in model parameters (***A*, Θ**) across EM iterations. (C) Mean GLM weight correlation between states per neuron. (D) Held-out log-likelihood for model trained on real and shuffled spikes from 132 target 2 reaches and tested on the 133^rd^ reach (Wilcoxon signed-rank test). Each dot represents the held-out trial.

We also checked how the differences in different GLM weight initializations with and without penalization impacted the final per-neuron GLM weights. Consistent with previous results from the mPFC dataset, ridge-regularized per-neuron initializations produced the most stable and biologically plausible final weights compared with unpenalized GLM or random weight initializations (Fig 6A). Lastly, we compared the convergence of the model using the trust-region optimization versus a standard Newton-Raphson method in the M-step (Fig 6B,C). Under Newton-Raphson, the negative log-likelihood increased across EM iterations, accompanied by over 50% non-finite GLM weights. In contrast, the trust-region method yielded more stable convergence with well-behaved finite parameter estimates. These results highlight the improved numerical stability of the trust-region approach.

**Figure 6.**
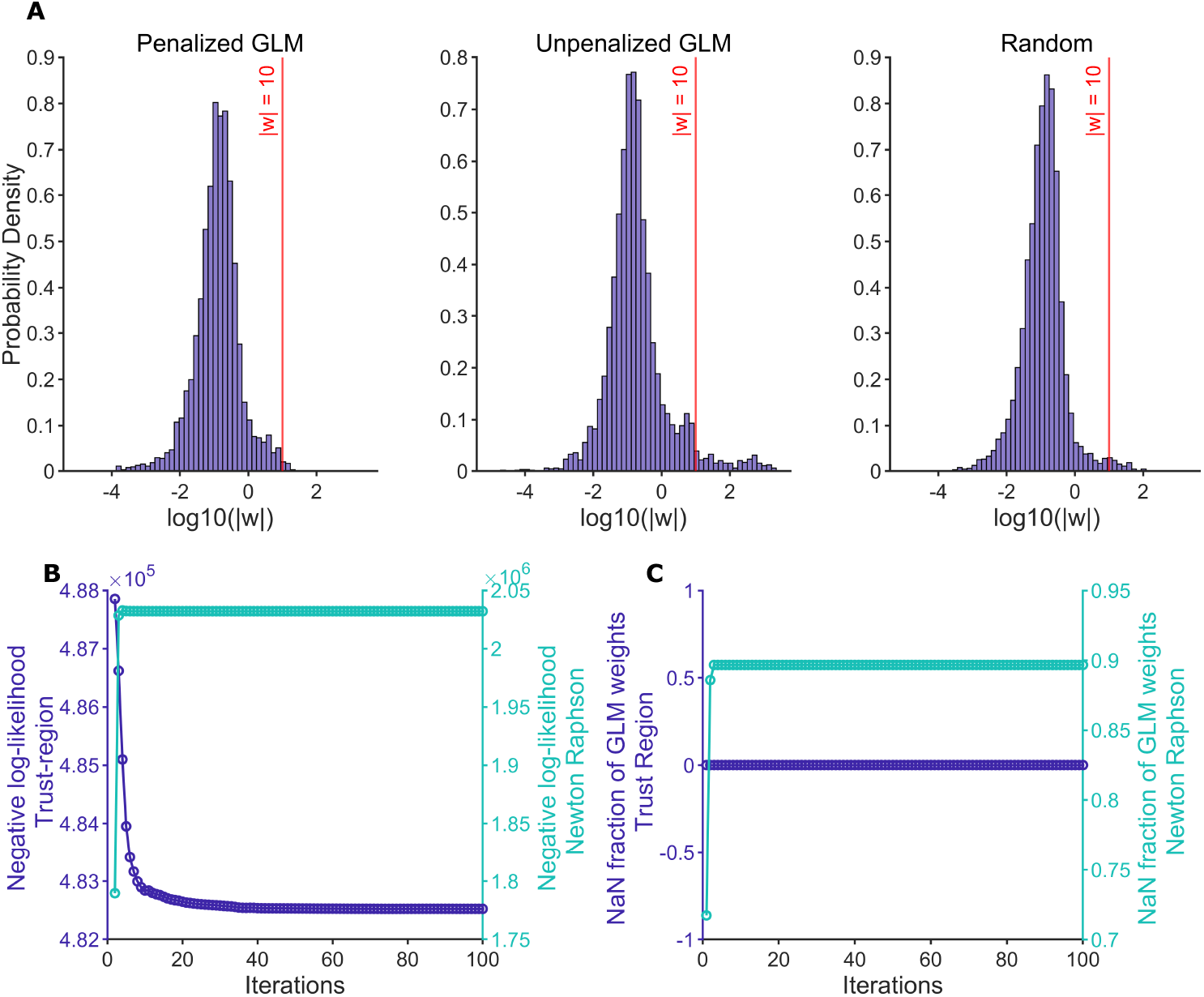
Initialization and optimization testing of the shared 2-state GLM-HMM fitted on primate premotor cortex activity. (A) Probability density distribution of final absolute GLM weights in log10 scale across all neurons using penalized GLM, unpenalized GLM, and random initializations. (B) Negative log-likelihood comparison across iterations when using trust-region and Newton-Raphson methods for M-step convergence. (C) Fraction of non-finite GLM weights across EM iterations when using trust-region and Newton-Raphson methods in the M-step.

### Anterior insula during alcohol self-administration

Lastly, we applied our methods to recordings made in the anterior insula while rats self-administered alcohol in a discrete-trial task (Supp. Fig 3). Specifically, each trial started with lever insertion, which served as a cue signaling alcohol availability, and the animal made a choice to seek or abstain from engaging with the lever to receive an alcohol reward (Fig 7A). We trained our model on a dataset comprising 45 seek and 27 abstain trials, with 96 simultaneously-recorded single units. Time-stamped events, including lever insertion, lever press, lever retraction, port entry, and spike history, were convolved with five raised-cosine basis functions over a 2.5-second time window. In addition to the temporal predictors, scalar predictors, including lever press latency and trial number, which were used to track the internal state, were also included in the predictor matrix. The data was binned in 50-millisecond time bins.

**Figure 7.**
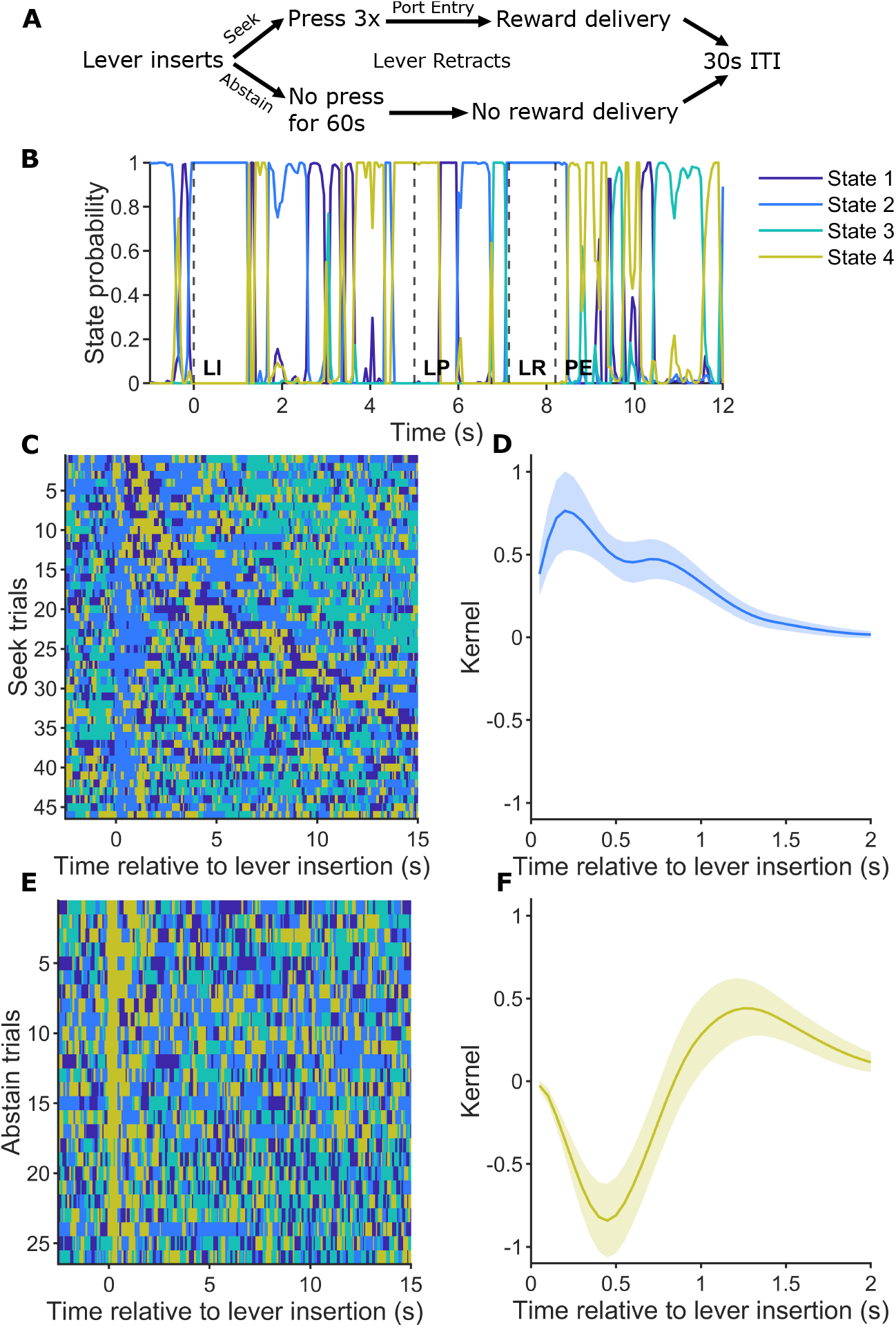
Distinct latent-state dynamics in the anterior insula during alcohol seeking and abstinence behavior. (A) Alcohol self-administration task schematic. (B) Posterior state probabilities during an example seek trial. (C) State probability heatmaps aligned to seek lever insertion (sorted by lever press latency). (D) Population-averaged kernel at seek lever insertion (Mean±SEM). (E) and (F) Same as (C) and (D) but aligned to abstain lever insertion.

Since we did not have prior knowledge about the true number of states, we tried different numbers of states, i.e., *S* ∈ {3, 4, 5}. The negative log-likelihood of the model fits showed a similar trend to the mPFC and premotor cortex dataset: as the number of states increased, the negative log-likelihoods decreased across EM iterations, and the distinction between states became subtle (Supp. Fig 4). Therefore, we decided to use a 4-state model.

The posterior probabilities for one seek trial are shown in Fig 7B, with state transitions around alcohol-associated cues (lever insertion and retraction) and alcohol intake (post-port entry). We compared the state probability heat maps for seek and abstain trials and observed that the state transitions in seek trials appeared more structured, when sorted by latency to lever press, than those in abstain trials (Fig 8C,E). Furthermore, the states observed at lever insertion for the two trial types are distinct: states 2 and 4 for seek and abstain trials, respectively. To understand the differences in ensemble activity across these states, we visualized the population-averaged kernels for states 2 and 4 (Fig 8D,F). The anterior insula population kernels showed a positive deviation for state 2 and a negative deviation for state 4. This corroborates previous findings of increased anterior insula engagement in the presence of alcohol cues to drive seeking behavior (Cofresí et al., 2019; Campbell et al., 2019).

**Figure 8.**
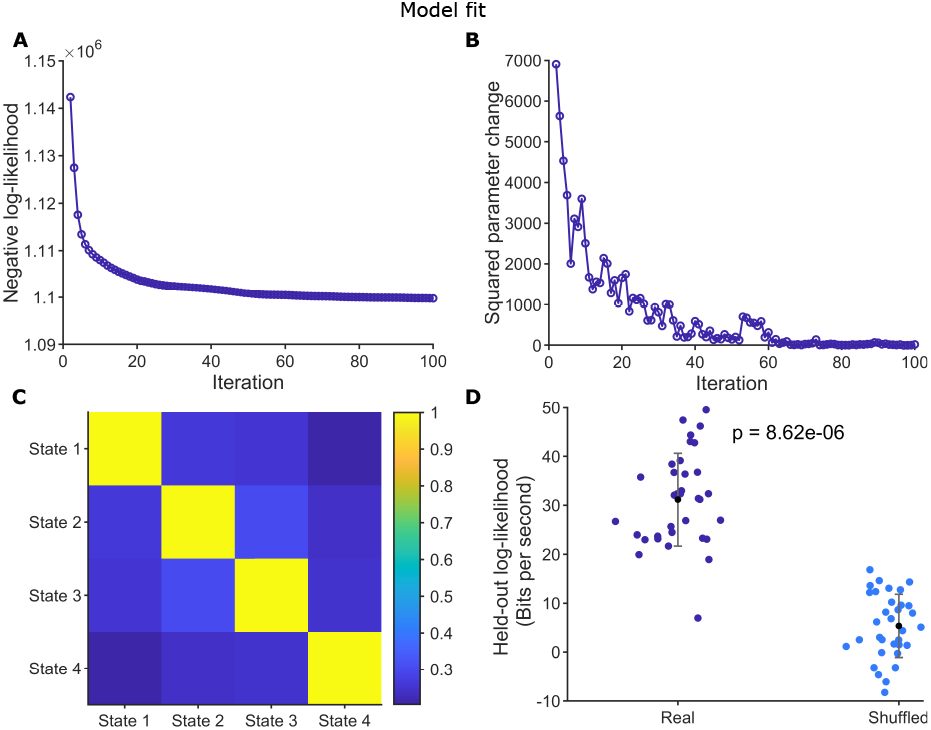
Convergence and validation of the 4-state GLM-HMM trained on anterior insula data. (A) Negative log-likelihood across EM iterations. (B) Change in model parameters (***A*, Θ**) across EM iterations. (C) Mean GLM weight correlation between states per neuron. (D) Held-out log-likelihood for model trained on real and shuffled spikes from 39 trials and tested on 40^th^ trial (Wilcoxon signed-rank test). Each dot represents the held-out trial.

To test our model fit, we tracked the convergence of our 4-state model’s negative log-likelihood and change in parameters across iterations (Fig 8A,B) and confirmed that the states were non-redundant (Fig 8C). We also tested the performance of the 4-state model trained on the first 40 trials and compared it with the model trained on shuffled spikes by computing the log-likelihood of observing the ensemble spiking in a held-out test trial. The log-likelihoods of the held-out test trials were significantly higher when the model was trained on the real dataset than when it was trained on a shuffled dataset (Fig 8D; Wilcoxon signed-rank test), highlighting that our model was able to select useful information when trained on nearly half the trials of the complete dataset.

When the 4-state model was initialized with a set of weights obtained by running a per-neuron ridge-penalized GLM and using that penalty in the M-step, as done for the above-mentioned datasets, the final GLM weights were well constrained within biological limits, whereas using a set of initial weights either by fitting a GLM with no penalization or using a random initialization yielded biologically implausible weights (Fig 9A). This underscores the importance of neuron-adaptive penalization for overcoming overfitting on sparse datasets. We also compared the M-step convergence of the dataset using the trust-region and Newton-Raphson methods. While the negative log-likelihood for the model decreased exponentially with the trust-region method, it increased exponentially with the Newton-Raphson method (Fig 9B). Furthermore, the Newton-Raphson method became increasingly unstable across iterations, as evidenced by the high fraction of non-finite GLM weights (Fig. 9C).

**Figure 9.**
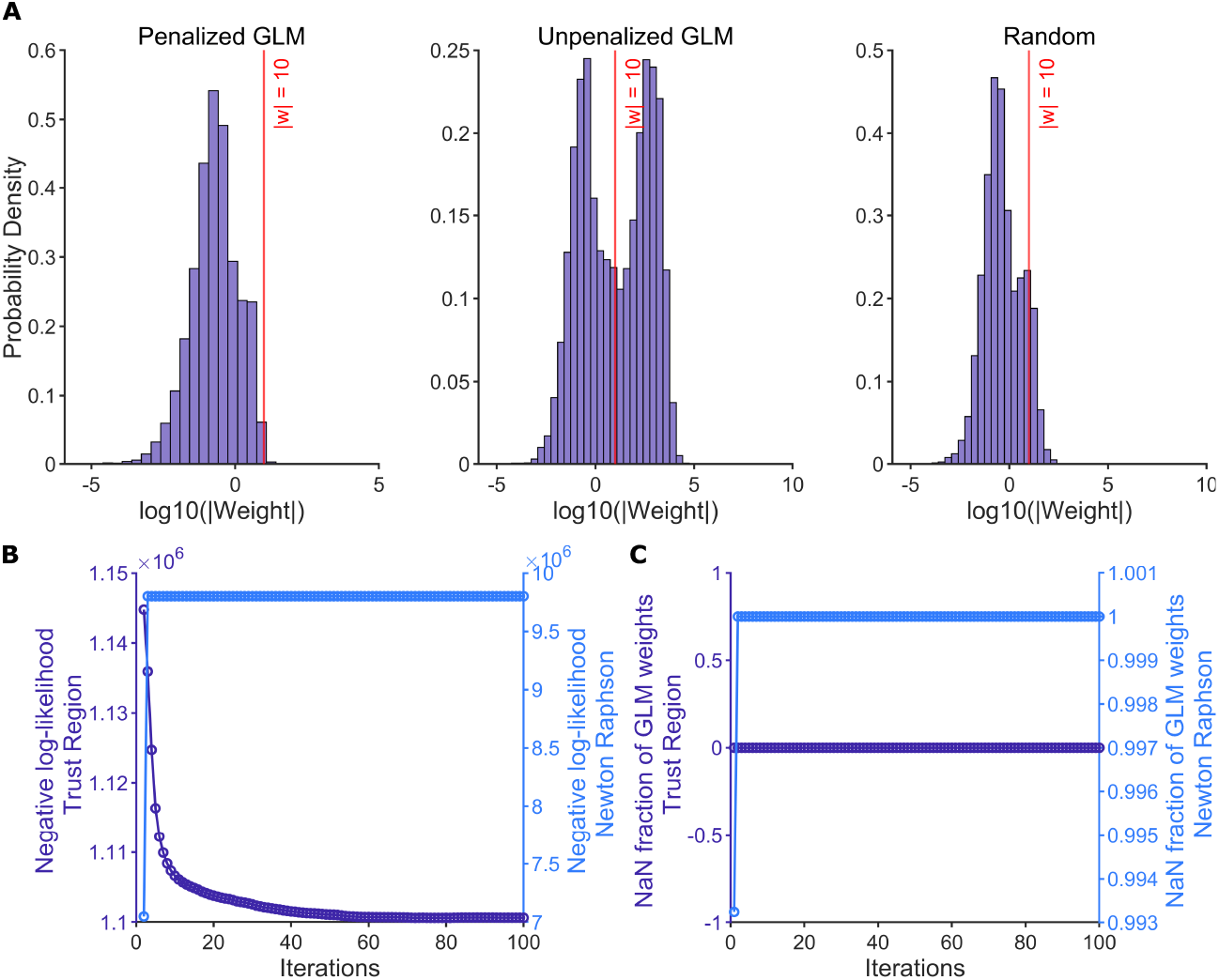
Comparison of initialization and optimization approaches for GLM-HMM fitted on rodent anterior insula activity. (A) Probability density distribution of final absolute GLM weights in log10 scale across all neurons using penalized GLM, unpenalized GLM, and random initializations. (B) Negative log-likelihood comparison across iterations when using trust-region and Newton-Raphson methods for M-step convergence. (C) Fraction of non-finite GLM weights across EM iterations when using trust-region and Newton-Raphson methods in the M-step.

## 6 Conclusion

In this work we developed and implemented a robust fitting procedure for multi-neuron GLM-HMM framework. Our work builds on prior work for neuronal GLM-HMMs (Escola et al., 2011), extending and adopting necessary changes to enable stable fitting on neuronal recordings. Our work addresses the key challenges that prevent current procedures used in, e.g., behavioral modeling with GLM-HMMs from working on neuronal data: the sparsity of neuronal activity, the limited data availability in many electrophysiology datasets, and the high correlations between stimulus features. Our framework incorporates the following features: 1) neuron-adaptive regularization in the M-step, obtained by fitting a five-fold cross-validated GLM to each neuron prior to model fitting, to yield biologically plausible GLM coefficients, 2) a trust-region optimization algorithm for stable M-step convergence when local curvature of the objective function is not quadratic, and 3) a leave-one-out trial cross-validation approach to test the model performance trained on limited data while preserving its temporal structure. Across three real electrophysiology datasets spanning multiple brain regions and species, as well as a series of benchmark analyses, our framework not only converges stably but also yields behaviorally interpretable latent states. In all, we have developed a practical approach for applying GLM-HMMs to neural population recordings, laying the groundwork for understanding how discrete latent states emerge from neuronal activity and shape behavior.

Our work complements the significant progress that has been made in identifying internal states by applying GLM-HMMs on behavioral datasets (Calhoun et al., 2019; Ashwood et al., 2022; Bolkan et al., 2022; Hulsey et al., 2024; Cuturela et al., 2025; Mohammadi et al., 2025; Li et al., 2023; Chen et al., 2026; Keeley et al., 2026; Moore et al., 2026). However, since internal states are an emergent property of neuronal population dynamics, models that directly incorporate ensemble spiking alongside task-related stimuli are needed to gain mechanistic insights underlying the latent states. In contrast to dynamical systems models that identify continuously evolving latent states (Sussillo et al., 2015; Kleinman et al., 2021; Pandarinath et al., 2018; Churchland et al., 2012; Mudrik et al., 2024; Yezerets et al., 2025; Mudrik et al., 2025), the categorical nature of HMMs in the GLM-HMM framework is often sought after to more directly model discrete behavioral modes. By removing the practical impediments of fitting GLM-HMM models to data, we aim to increase the ability of neuroscientists to study such time-varying neural activity.

## Acknowledgements

This work was supported by the National Institutes of Health R01AA031609, R01AA027213, and U01NS115587 grants, and NSF CAREER Award 2340338. We are incredibly thankful to Dr. Timothy Harris and his student, Sarah Jung, for facilitating Neuropixels 2.0 recordings in the anterior insula.

## Data availability

The prefrontal cortex and premotor cortex data are both publicly available datasets (Peyrache et al., 2009, 2018; Lawlor et al., 2018; Perich et al., 2018b). Data for the anterior insula example is available as part of the Github database at https://github.com/AamnaLawrence/GLM-HMM-Neuronal-Encoding/tree/main, with Zenodo DOI https://doi.org/10.5281/zenodo.20970583.

## Code availability

The MATLAB implementation of our GLM-HMM fitting is available on Github at https://github.com/AamnaLawrence/GLM-HMM-Neuronal-Encoding/tree/main, with Zenodo DOI https://doi.org/10.5281/zenodo.20970583.

## Supplementary Information

### Scaling and normalization of EM probabilities

The forward probabilities computed in Equation (22) are summed for all neurons and normalized by Σ_*s*_ *α*_*ts*_. Thus, the final forward probabilities in log-space and after scaling are,

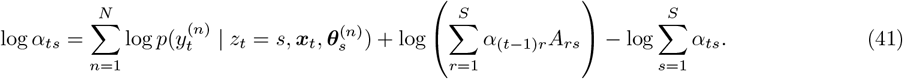

Similarly, Equation (25) after scaling with Σ_*r*_ *β*_*tr*_ is given by,

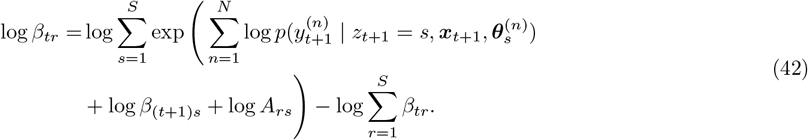

Equation (16) in log-space with normalization is written as,

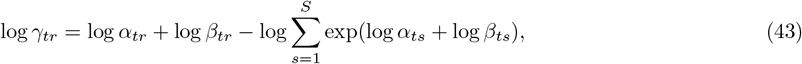

and Equation (28) transforms into,

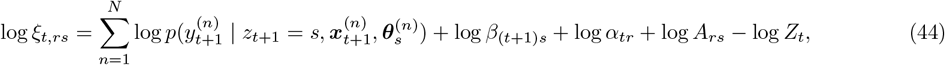

where

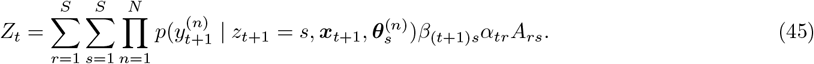

### mPFC dataset

As noted in the main text, we tested our model on the mPFC dataset for *S* ∈ {3, 4, 5} states. Across 100 EM iterations, our model showed a decreasing negative log-likelihood as the state number increased (Supp. Fig 1A,B,C). However, the differences between the states got subtle based on the average per-neuron GLM weight correlations (Supp. Fig 1D,E,F).

### Premotor cortex dataset

Similar to the mPFC dataset, we tested our model on the premotor cortex dataset for *S* ∈ {2, 3} states. Over 100 EM iterations, our model showed a decreasing negative log-likelihood as the number of states increased (Supp. Fig 2A,B). However, the differences between the states got subtle based on the average per-neuron GLM weight correlations (Supp. Fig 2C,D).

### Anterior insula dataset

The anterior insula dataset was collected from a rat trained to self-administer alcohol (Supp. Fig 3A). The rat was given intermittent pre-exposure to 15% EtOH in their home cage three times a week, 24 hours at a time for a period of 5 weeks prior to operant training. This was conducted to acclimatize them to the taste and pharmacological effects of ethanol and allow stabilization of drinking behavior (≥ 2g/kg in the last week of the drinking sessions) prior to instrumental training.

On each trial, the rat was trained to press a lever three times to earn a bolus of 10% alcohol reward, where each trial began with a lever insertion. Three responses on the lever within 60 seconds of lever insertion triggered lever retraction and reward delivery, followed by a 30-second inter-trial interval. These sessions were limited to 1 hour or 30 reward deliveries, whichever occurred first. After stable behavioral performance, the rat underwent chronic Neuropixels 2.0 implantation in the anterior insula (see Supp. Fig 3B for recording site verification) and was retrained. Recordings were performed. All data were acquired with SpikeGLX and hand-sorted with Kilosort 4.0.

To determine the optimal number of states, we tested our model on the anterior insula dataset for *S* ∈ {3, 4, 5} states. Across 100 EM iterations, our model showed a decreasing negative log-likelihood as the number of states increased (Supp. Fig 4A,B,C). However, the states got more correlated based on the average per-neuron GLM weight correlations (Supp. Fig 4D,E,F).

**Supplementary Figure 1.**
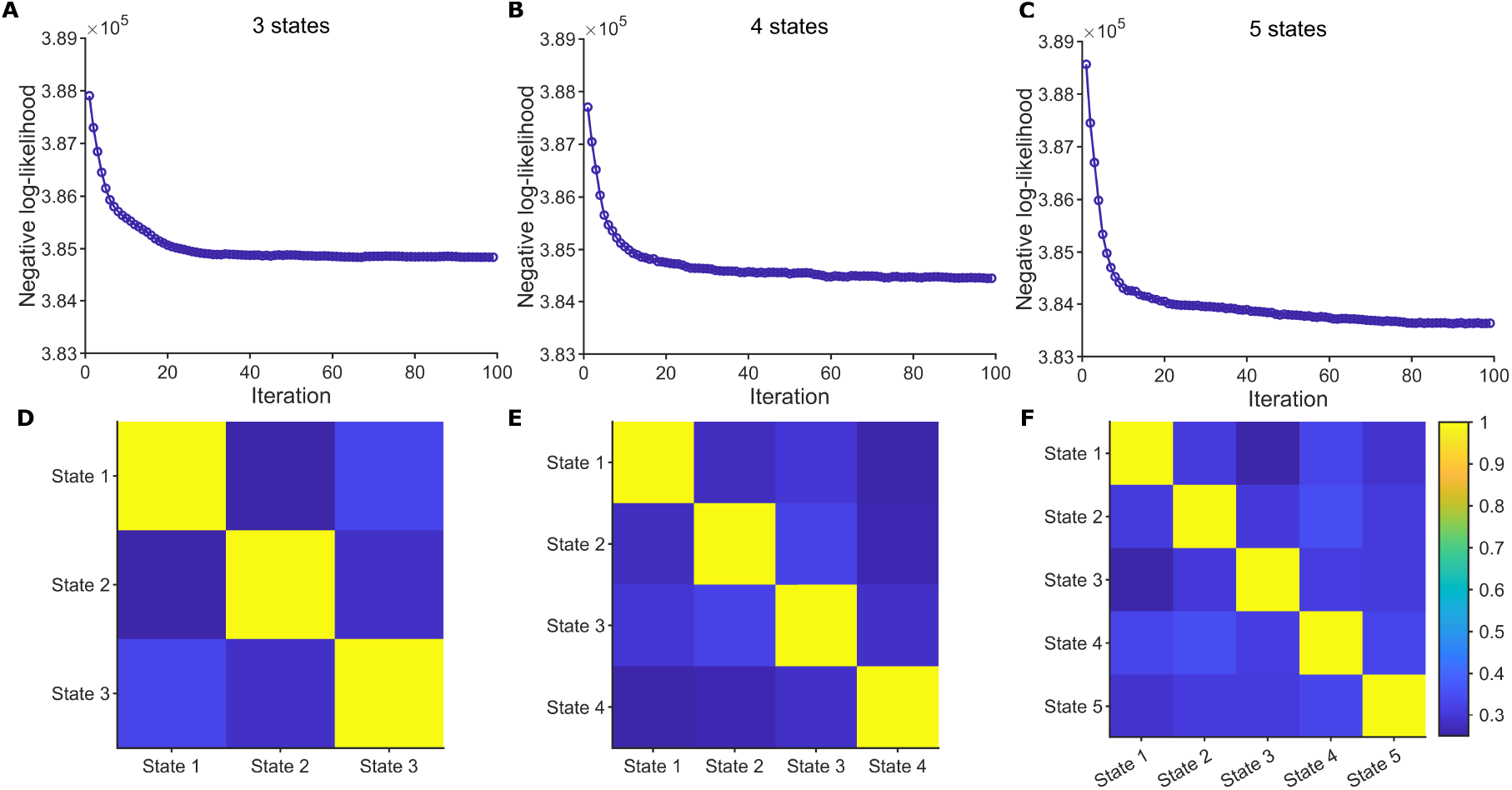
Evaluation of a shared 3-state, 4-state, and 5-state GLM-HMM trained on rodent mPFC dataset. Negative log-likelihood across 100 EM iterations for (A) 3-state, (B) 4-state, (C) 5-state model. Mean GLM weight correlation between states per neuron for (D) 3-state, (E) 4-state, (F) 5-state model.

**Supplementary Figure 2.**
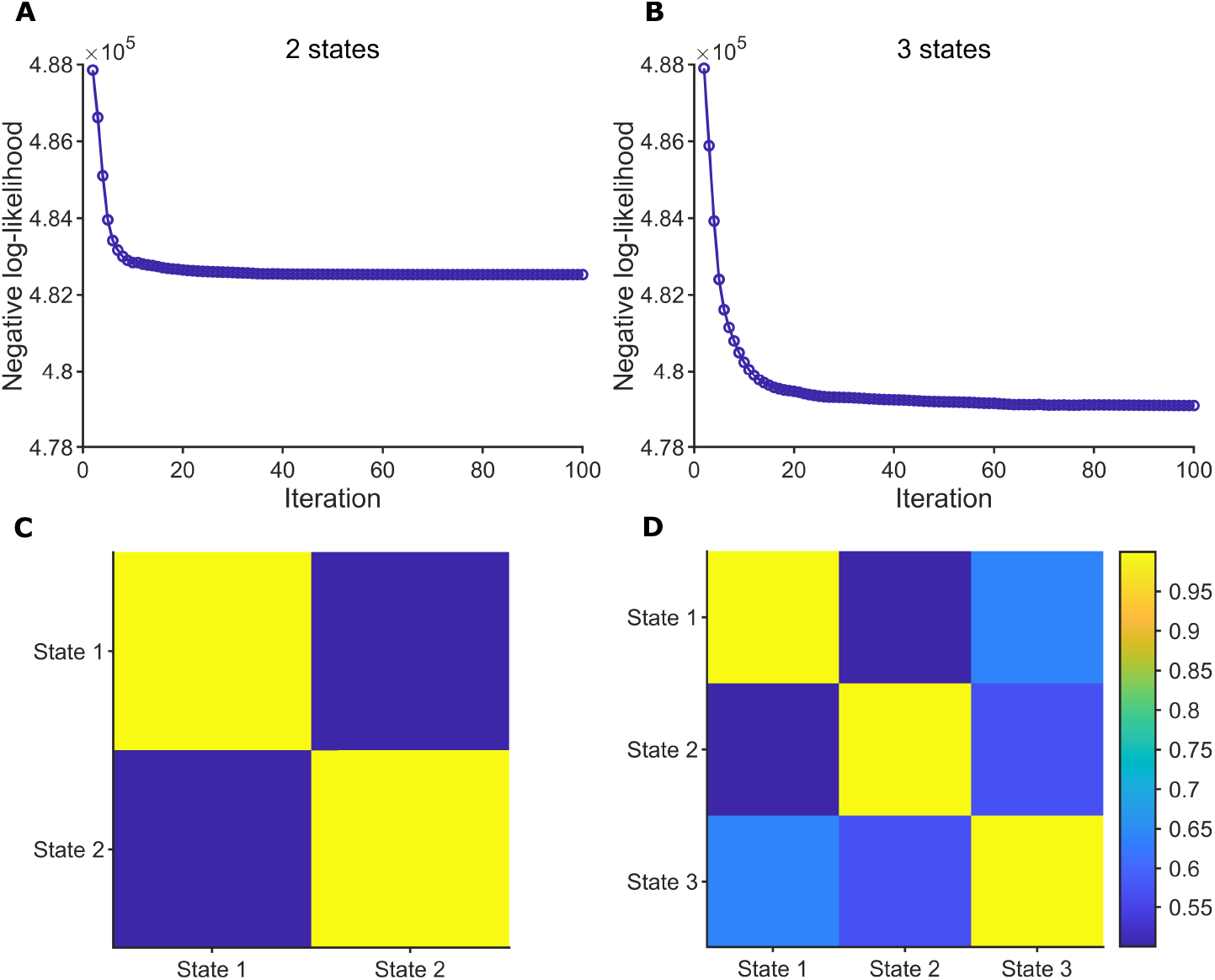
Evaluation of a shared 2-state and 3-state GLM-HMM trained on primate premotor cortex dataset. Negative log-likelihood across 100 EM iterations for (A) 2-state, (B) 3-state model. Mean GLM weight correlation between states per neuron for (C) 2-state, (D) 3-state model.

**Supplementary Figure 3.**
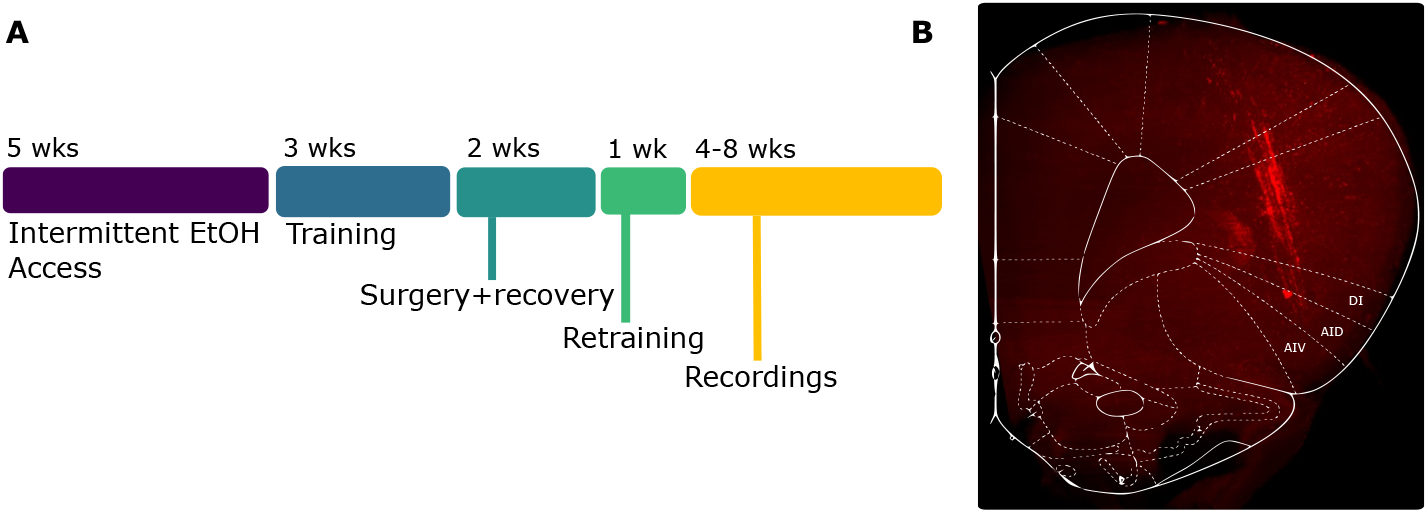
Experimental procedure and histological validation of recording location. (A) Experimental timeline. (B) Cleared rodent brain image showing DiI-coated Neuropixels 2.0 probe shank trajectories in the anterior insula (AIV: agranular insula ventral; AID: agranular insula dorsal; DI: dysgranular insula).

**Supplementary Figure 4.**
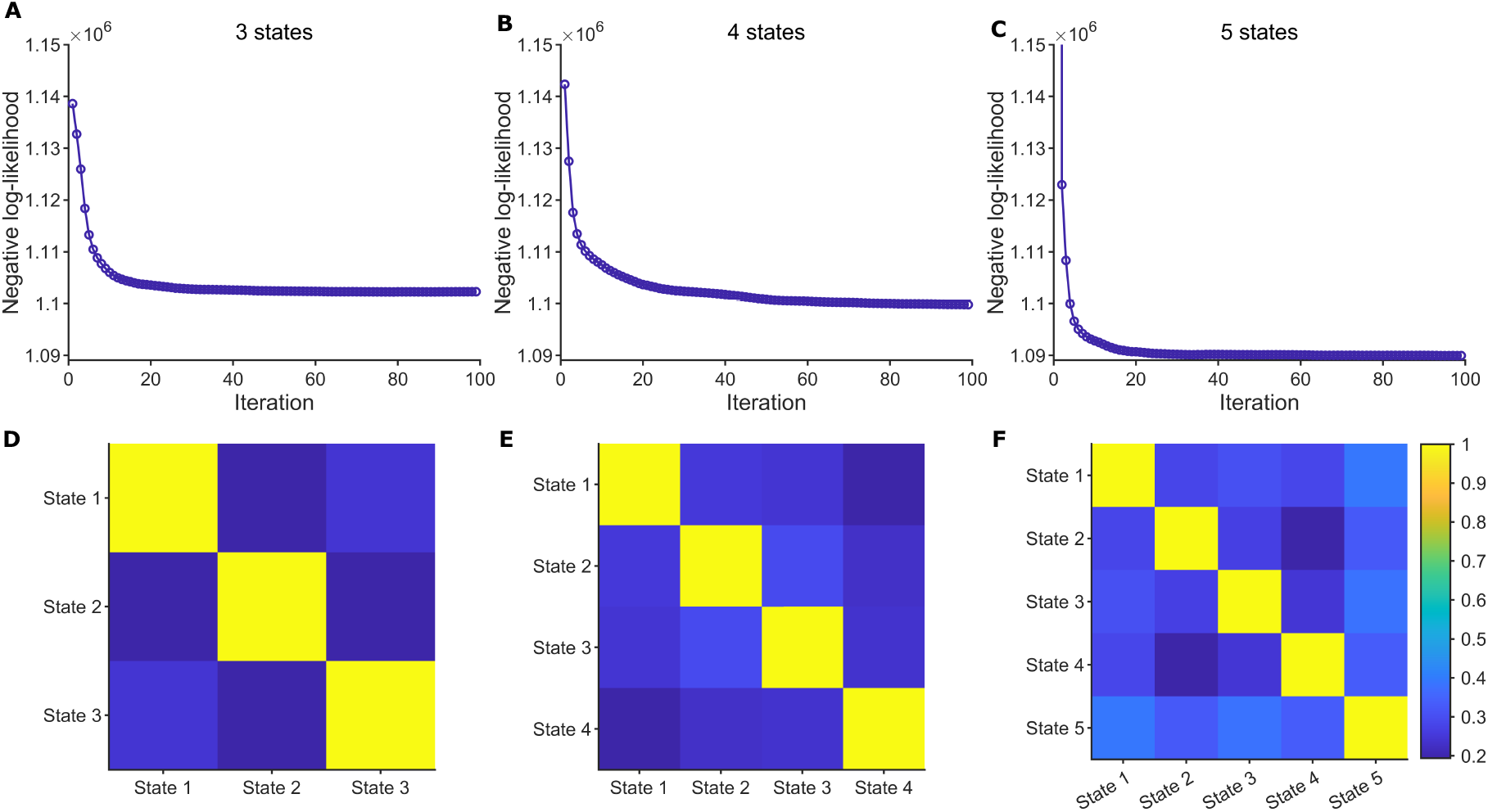
Evaluation of a shared 3-state, 4-state, and 5-state GLM-HMM trained on rodent anterior insula dataset. Negative log-likelihood across 100 EM iterations for (A) 3-state, (B) 4-state, (C) 5-state model. Mean GLM weight correlation between states per neuron for (D) 3-state, (E) 4-state, (F) 5-state model.

## References

David J. Anderson. Circuit modules linking internal states and social behaviour in flies and mice. Nature Reviews Neuroscience, 17(11):692–704, 2016. doi: 10.1038/nrn.2016.125. URL https://doi.org/10.1038/nrn.2016.125.

Yvan Aponte, Duygu Atasoy, and Scott M. Sternson. Agrp neurons are sufficient to orchestrate feeding behavior rapidly and without training. Nature Neuroscience, 14(3):351–355, 2011. doi: 10.1038/nn.2739. URL https://doi.org/10.1038/nn.2739.

Kazunori Asahina, Kimiko Watanabe, Barbara J. Duistermars, Eric Hoopfer, Carlos R. González, Erla A. Eyjólfsdóttir, Pietro Perona, and David J. Anderson. Tachykinin-expressing neurons control male-specific aggressive arousal in drosophila. Cell, 156(1–2):221–235, 2014. doi: 10.1016/j.cell.2013.11.045. URL https://doi.org/10.1016/j.cell.2013.11.045.

Zoe C. Ashwood, Nicholas A. Roy, Ian R. Stone, Anne E. Urai, Anne K. Churchland, and Jonathan W. Pillow. Mice alternate between discrete strategies during perceptual decision-making. Nature Neuroscience, 25(2):201–212, 2022. doi: 10.1038/s41593-021-01007-1.

Leonard E. Baum, Ted Petrie, George Soules, and Norman Weiss. A maximization technique occurring in the statistical analysis of probabilistic functions of markov chains. The Annals of Mathematical Statistics, 41(1): 164–171, 1970. doi: 10.1214/aoms/1177697196.

Yoshua Bengio and Paolo Frasconi. An input output HMM architecture. In Gerald Tesauro, David S. Touretzky, and Todd K. Leen, editors, Advances in Neural Information Processing Systems 7 (NIPS 1994), pages 427–434. MIT Press, 1995.

Jorge Berrios, Chen Li, Joseph C. Madara, Alastair S. Garfield, J. Scott Steger, Michael J. Krashes, and Bradford B. Lowell. Food cue regulation of agrp hunger neurons guides learning. Nature, 595(7866):695–700, 2021. doi: 10.1038/s41586-021-03729-3. URL https://doi.org/10.1038/s41586-021-03729-3.

Jennifer M. Birrell and Verity J. Brown. Medial frontal cortex mediates perceptual attentional set shifting in the rat. Journal of Neuroscience, 20(11):4320–4324, 2000. ISSN 0270-6474. doi: 10.1523/JNEUROSCI.20-11-04320.2000. URL https://www.jneurosci.org/content/20/11/4320.

Steven S. Bolkan, Joshua M. Stujenske, Sébastien Parnaudeau, Timothy J. Spellman, Claire Rauffenbart, Abdullah I. Abbas, Adam Z. Harris, and Joshua A. Gordon. Opponent control of behavior by dorsomedial striatal pathways depends on task demands and internal state. Nature Neuroscience, 25(5):651–661, 2022. doi: 10.1038/s41593-022-01022-4.

Adam J. Calhoun, Jonathan W. Pillow, and Mala Murthy. Unsupervised identification of the internal states that shape natural behavior. Nature Neuroscience, 22(12):2040–2049, 2019. doi: 10.1038/s41593-019-0533-x.

Emma J. Campbell, Jennifer P. M. Flanagan, Laura C. Walker, Matthew K. R. I. Hill, Nathan J. Marchant, and Andrew J. Lawrence. Anterior insular cortex is critical for the propensity to relapse following punishment-imposed abstinence of alcohol seeking. Journal of Neuroscience, 39(6):1077–1087, 2019. doi: 10.1523/JNEUROSCI.1596-18.2018. URL https://doi.org/10.1523/JNEUROSCI.1596-18.2018.

Kevin S. Chen, Jonathan W. Pillow, and Andrew M. Leifer. State-switching navigation strategies in Caenorhabditis elegans are beneficial for chemotaxis. Proceedings of the National Academy of Sciences, 123:e2519999123, 2026. doi: 10.1073/pnas.2519999123.

Yu Chen, Ryan A. Essner, Sercan Kosar, Olivia H. Miller, Yi-Chun Lin, Sanaz Mesgarzadeh, and Zachary A. Knight. Sustained npy signaling enables agrp neurons to drive feeding. eLife, 8:e46348, 2019. doi: 10.7554/eLife.46348. URL https://doi.org/10.7554/eLife.46348.

Mark M. Churchland, John P. Cunningham, Matthew T. Kaufman, John D. Foster, Paul Nuyujukian, Stephen I. Ryu, and Krishna V. Shenoy. Neural population dynamics during reaching. Nature, 487(7405):51–56, 2012. doi: 10.1038/nature11129. URL https://doi.org/10.1038/nature11129.

Matthew A. Churgin, Robert J. McCloskey, Emily Peters, and Christopher Fang-Yen. Antagonistic serotonergic and octopaminergic neural circuits mediate food-dependent locomotory behavior in caenorhabditis elegans. Journal of Neuroscience, 37(32):7811–7823, 2017. doi: 10.1523/JNEUROSCI.2636-16.2017. URL https://doi.org/10.1523/JNEUROSCI.2636-16.2017.

Emily J. Clowney, Shinya Iguchi, Joshua J. Bussell, Edward Scheer, and Vanessa Ruta. Multimodal chemosensory circuits controlling male courtship in drosophila. Neuron, 87(5):1036–1049, 2015. doi: 10.1016/j.neuron.2015.07.025. URL https://doi.org/10.1016/j.neuron.2015.07.025.

Roberto U. Cofresí, Dylan J. Grote, Eric Viet Thanh Le, Marie-H. Monfils, Nadia Chaudhri, Rueben A. Gonzales, and Hongjoo J. Lee. Alcohol-associated antecedent stimuli elicit alcohol seeking in non-dependent rats and may activate the insula. Alcohol, 76:91–102, 2019. ISSN 0741-8329. doi: 10.1016/j.alcohol.2018.08.004. URL https://doi.org/10.1016/j.alcohol.2018.08.004.

L. I. Cuturela, International Brain Laboratory, and Jonathan W. Pillow. Internal states emerge early during learning of a perceptual decision-making task. bioRxiv, 2025. doi: 10.1101/2024.11.30.626182. URL https://www.biorxiv.org/content/10.1101/2024.11.30.626182v1. Preprint. PMID: 39651276; PMCID: PMC11623682.

Arthur P. Dempster, Nan M. Laird, and Donald B. Rubin. Maximum likelihood from incomplete data via the em algorithm. Journal of the Royal Statistical Society: Series B (Methodological), 39(1):1–38, 1977. doi: 10.1111/j. 2517-6161.1977.tb01600.x.

Sean Escola, Alfredo Fontanini, Don Katz, and Liam Paninski. Hidden markov models for the stimulus-response relationships of multistate neural systems. Neural computation, 23(5):1071–1132, 2011.

Steven W. Flavell, Nikhil Pokala, Evan Z. Macosko, Dirk R. Albrecht, Jennifer Larsch, and Cornelia I. Bargmann. Serotonin and the neuropeptide pdf initiate and extend opposing behavioral states in c. elegans. Cell, 154(5): 1023–1035, 2013. doi: 10.1016/j.cell.2013.08.001. URL https://doi.org/10.1016/j.cell.2013.08.001.

Stan B. Floresco, David N. Braaksma, and Anthony G. Phillips. Thalamic-cortical-striatal circuitry subserves working memory during delayed responding on a radial arm maze. Journal of Neuroscience, 19(24):11061–11071, 1999. doi: 10.1523/JNEUROSCI.19-24-11061.1999.

Joshua I. Gold and Michael N. Shadlen. The neural basis of decision making. Annual Review of Neuroscience, 30: 535–574, 2007. doi: 10.1146/annurev.neuro.29.051605.113038.

David M. Green and John A. Swets. Signal Detection Theory and Psychophysics. Wiley, New York, 1966.

Jan Gründemann, Yael Bitterman, Tingting Lu, Steffen Krabbe, Benjamin F. Grewe, Mark J. Schnitzer, and Andreas Lüthi. Amygdala ensembles encode behavioral states. Science, 364(6437):eaav8736, 2019. doi: 10.1126/science.aav8736. URL https://doi.org/10.1126/science.aav8736.

Tomas Hindmarsh Sten, Rui Li, Anne Otopalik, and Vanessa Ruta. Sexual arousal gates visual processing during drosophila courtship. Nature, 595(7866):549–553, 2021. doi: 10.1038/s41586-021-03714-w. URL https://doi.org/10.1038/s41586-021-03714-w.

Daniel R. Hulsey, Katherine Zumwalt, Luca Mazzucato, et al. Decision-making dynamics are predicted by arousal and uninstructed movements. Cell Reports, 43(2), 2024.

Na Ji, Gaurav K. Madan, Guillaume I. Fabre, Adam Dayan, Cody M. Baker, Tyler S. Kramer, Ifeoma Nwabudike, and Steven W. Flavell. A neural circuit for flexible control of persistent behavioral states. eLife, 10:e62889, 2021. doi: 10.7554/eLife.62889. URL https://doi.org/10.7554/eLife.62889.

Matthew W. Jones and Michael A. Wilson. Theta rhythms coordinate hippocampal–prefrontal interactions in a spatial memory task. PLOS Biology, 3(12):e402, 2005. doi: 10.1371/journal.pbio.0030402.

Tomomi Karigo and Adam S. Charles. Towards a multi-dimensional understanding of brain states. Neurobiology of Learning and Memory, 222:108110, 2025. ISSN 1074-7427. doi: 10.1016/j.nlm.2025.108110. URL https://www.sciencedirect.com/science/article/pii/S1074742725000917.

Tomomi Karigo, Adam Kennedy, Bin Yang, Yeonhee Kim, Marc Tessier-Lavigne, and David J. Anderson. Distinct hypothalamic control of same- and opposite-sex mounting behaviour in mice. Nature, 589(7841):258–263, 2021. doi: 10.1038/s41586-020-2995-0. URL https://doi.org/10.1038/s41586-020-2995-0.

Stephen Keeley, David Zoltowski, Adam Charles, and Jonathan Pillow. Improved inference of latent neural states from calcium imaging data. eLife, 15, 2026.

Michael Kleinman, Chandramouli Chandrasekaran, and Jonathan Kao. A mechanistic multi-area recurrent network model of decision-making. In Marc’Aurelio Ranzato, Alina Beygelzimer, Yann Dauphin, Percy S. Liang, and Jennifer Wortman Vaughan, editors, Advances in Neural Information Processing Systems, volume 34, pages 23152–23165. Curran Associates, Inc., 2021.

Patrick N. Lawlor, Matthew G. Perich, Lee E. Miller, and Konrad P. Kording. Linear-nonlinear-time-warp-poisson models of neural activity. Journal of Computational Neuroscience, 45(3):173–191, 2018. doi: 10.1007/s10827-018-0696-6.

Chengrui Li, Soon Ho Kim, Chris Rodgers, Hannah Choi, and Anqi Wu. One-hot generalized linear model for switching brain state discovery, 2023. URL https://arxiv.org/abs/2310.15263.

Yonatan Livneh, Ramakrishnan N. Ramesh, Christopher R. Burgess, Kevin M. Levandowski, Joseph C. Madara, Helen Fenselau, Griffin J. Goldey, Veronica E. Diaz, Nick Jikomes, Jacob M. Resch, Bradford B. Lowell, and Mark L. Andermann. Homeostatic circuits selectively gate food cue responses in insular cortex. Nature, 546(7660): 611–616, 2017. doi: 10.1038/nature22375. URL https://doi.org/10.1038/nature22375.

Yonatan Livneh, Alexander U. Sugden, Joseph C. Madara, Ryan A. Essner, Victor I. Flores, Lauren A. Sugden, Jacob M. Resch, Bradford B. Lowell, and Mark L. Andermann. Estimation of current and future physiological states in insular cortex. Neuron, 105(6):1094–1111.e10, 2020. doi: 10.1016/j.neuron.2019.12.027. URL https://doi.org/10.1016/j.neuron.2019.12.027.

João C. Marques, Miao Li, Daniel Schaak, Drew N. Robson, and Jing M. Li. Internal state dynamics shape brainwide activity and foraging behaviour. Nature, 577(7789):239–243, 2020. doi: 10.1038/s41586-019-1858-z. URL https://doi.org/10.1038/s41586-019-1858-z.

Alexander Mathis, Pranav Mamidanna, Kevin M. Cury, Taiga Abe, Venkatesh N. Murthy, Mackenzie W. Mathis, and Matthias Bethge. DeepLabCut: markerless pose estimation of user-defined body parts with deep learning. Nature Neuroscience, 21(9):1281–1289, 2018. doi: 10.1038/s41593-018-0209-y. URL https://doi.org/10.1038/s41593-018-0209-y.

Arina Medvedeva, Edoardo Balzani, Alex H. Williams, and Stephen L. Keeley. Scalable inference of functional neural connectivity at submillisecond timescales, 2025. URL 10.48550/arXiv.2510.20966.

Z. Mohammadi, Zoe C. Ashwood, The International Brain Laboratory, et al. Identifying the factors governing internal state switches during nonstationary sensory decision-making. Nature Communications, 16:11684, 2025. doi: 10.1038/s41467-025-66738-0. URL https://doi.org/10.1038/s41467-025-66738-0.

Sharlen Moore, Zyan Wang, Ziyi Zhu, Joy Wang, Ruolan Sun, Yeonjae A Lee, Adam Charles, and Kishore V Kuchibhotla. Revealing abrupt transitions from goal-directed to habitual behavior. Nature Communications, 2026.

Noga Mudrik, Yenho Chen, Eva Yezerets, Christopher J Rozell, and Adam S Charles. Decomposed Linear Dynamical Systems (dLDS) for learning the latent components of neural dynamics. Journal of Machine Learning Research, 25:1–12, 2024.

Noga Mudrik, Ryan Ly, Oliver Ruebel, and Adam S Charles. Creimbo: Cross-regional ensemble interactions in multi-view brain observations. The International Conference on Learning Representations, 2025.

Chethan Pandarinath, Daniel J. O’Shea, Jasmine Collins, Rafal Jozefowicz, Sergey D. Stavisky, Jonathan C. Kao, Eric M. Trautmann, Matthew T. Kaufman, Stephen I. Ryu, Leigh R. Hochberg, Jaimie M. Henderson, and Krishna V. Shenoy. Inferring single-trial neural population dynamics using sequential auto-encoders. Nature Methods, 15(10):805–815, 2018. doi: 10.1038/s41592-018-0109-9. URL https://doi.org/10.1038/s41592-018-0109-9.

Liam Paninski. Maximum likelihood estimation of cascade point-process neural encoding models. Network: Computation in Neural Systems, 15(4):243–262, 2004. doi: 10.1088/0954-898X/15/4/002. URL https://doi.org/10.1088/0954-898X/15/4/002.

Il Memming Park, Miriam L. R. Meister, Alexander C. Huk, and Jonathan W. Pillow. Encoding and decoding in parietal cortex during sensorimotor decision-making. Nature Neuroscience, 17(10):1395–1403, 2014. doi: 10.1038/nn.3800. URL https://doi.org/10.1038/nn.3800.

Talmo D. Pereira, Nathan Tabris, Arie Matsliah, Daniel M. Turner, Junyu Li, Sridhar Ravindranath, Elias S. Papadoyannis, Emmanuel Normand, David S. Deutsch, Z. Yujia Wang, Grace C. McKenzie-Smith, Catalin C. Mitelut, Luisa N. Castro, Jamie D’Uva, Mikhail Kislin, Joshua R. Sanes, Sarah D. Kocher, Shaul Wang, Annegret L. Falkner, Joshua W. Shaevitz, and Mala Murthy. SLEAP: A deep learning system for multi-animal pose tracking. Nature Methods, 19(4):486–495, 2022. doi: 10.1038/s41592-022-01426-1. URL https://doi.org/10.1038/s41592-022-01426-1.

Matthew G. Perich, Patrick N. Lawlor, Konrad P. Kording, and Lee E. Miller. Extracellular neural recordings from macaque primary and dorsal premotor motor cortex during a sequential reaching task, 2018a. URL 10.6080/K0FT8J72.

Matthew G. Perich, Patrick N. Lawlor, Konrad P. Kording, and Lee E. Miller. Extracellular neural recordings from macaque primary and dorsal premotor motor cortex during a sequential reaching task, 2018b. URL 10.6080/K0FT8J72.

Adrien Peyrache, Mehdi Khamassi, Karim Benchenane, Samuel I. Wiener, and Francesco P. Battaglia. Replay of rule-learning related neural patterns in the prefrontal cortex during sleep. Nature Neuroscience, 12(7):919–926, 2009. doi: 10.1038/nn.2337.

Adrien Peyrache, Mehdi Khamassi, Karim Benchenane, Sidney I. Wiener, and Francesco P. Battaglia. Activity of neurons in rat medial prefrontal cortex during learning and sleep, 2018. URL 10.6080/K0KH0KH5.

Lawrence R. Rabiner. A tutorial on hidden markov models and selected applications in speech recognition. Proceedings of the IEEE, 77(2):257–286, 1989. doi: 10.1109/5.18626. URL https://doi.org/10.1109/5.18626.

Roger Ratcliff and Gail McKoon. The diffusion decision model: Theory and data for two-choice decision tasks. Neural Computation, 20(4):873–922, 2008. doi: 10.1162/neco.2008.12-06-420.

Roger Ratcliff and Jeffrey N. Rouder. Modeling response times for two-choice decisions. Psychological Science, 9(5): 347–356, 1998. doi: 10.1111/1467-9280.00067.

Eero P. Simoncelli, Liam Paninski, Jonathan W. Pillow, and Odelia Schwartz. Characterization of neural responses with stochastic stimuli. In Michael S. Gazzaniga, editor, The Cognitive Neurosciences, pages 327–338. MIT Press, Cambridge, MA, 3 edition, 2004.

David Sussillo, Mark M. Churchland, Matthew T. Kaufman, and Krishna V. Shenoy. A neural network that finds a naturalistic solution for the production of muscle activity. Nature Neuroscience, 18(7):1025–1033, 2015. doi: 10.1038/nn.4042. URL https://doi.org/10.1038/nn.4042.

Wilson Truccolo, Uri T. Eden, Matthew R. Fellows, John P. Donoghue, and Emery N. Brown. A point process framework for relating neural spiking activity to spiking history, neural ensemble, and extrinsic covariate effects. Journal of Neurophysiology, 93(2):1074–1089, 2005. doi: 10.1152/jn.00697.2004. URL https://doi.org/10.1152/jn.00697.2004.

Anqi Wu, Estefany Kelly Buchanan, Matthew Whiteway, Michael Schartner, Guido Meijer, Jean-Paul Noel, Erica Rodriguez, Claire Everett, Amy Norovich, Evan Schaffer, Neeli Mishra, C. Daniel Salzman, Dora Angelaki, Andrés Bendesky, The International Brain Laboratory, John Cunningham, and Liam Paninski. Deep graph pose: a semi-supervised deep graphical model for improved animal pose tracking. In Hugo Larochelle, Marc’Aurelio Ranzato, Raia Hadsell, Maria-Florina Balcan, and Hsuan-Tien Lin, editors, Advances in Neural Information Processing Systems, volume 33, pages 6040–6052. Curran Associates, Inc., 2020.

Eva Yezerets, Noga Mudrik, and Adam S CharlesDecomposed linear dynamical systems (dlds) models reveal instantaneous, context-dependent dynamic connectivity in c. elegans. Communications biology, 8(1):1218, 2025.

